# Biological landscape of acute illness in children in sub-Saharan Africa and South Asia

**DOI:** 10.1101/2025.09.01.672585

**Authors:** Evans O. Mudibo, Charles Sande, Moses M Ngari, Benjamin Jenkins, Benson O. Singa, Christina L. Lancioni, Abdoulaye Hama Diallo, Emmie Mbale, Ezekiel Mupere, John Mukisa, Roseline Maïmouna Bamouni, Andrew J. Prendergast, Christine J. McGrath, Albert Koulman, Robert H.J. Bandsma, Kirkby D. Tickell, Judd L Walson, James A. Berkley, Gerard Bryan Gonzales, James M. Njunge

## Abstract

Childhood illnesses including pneumonia, diarrhoea and malaria are leading causes of hospitalisation and mortality in resource-limited settings. However, we lack understanding of whether systemic responses to such diverse clinical syndromes are shared or specific, how they are impacted by malnutrition and how they differ from well children. We performed multi-omic profiling of plasma proteins, and serum metabolites and lipids in acutely ill hospitalised and well children in sub-Saharan Africa and South Asia. Using network-based clustering and mixed-effects modelling, we identified common and syndrome-specific omics responses to acute illness. We found that malnutrition often modifies host responses to disease. Although the internal structure of individual omics modules was largely preserved between ill and well children, the interactions between these preserved modules were markedly reorganised during acute illness. Compared to well children, biological systems in hospitalised children were more interconnected, exhibiting denser cross-omics interactions. These findings reveal widespread multisystem mobilisation during paediatric acute illness, offer deeper mechanistic insights and highlight candidate pathways for therapeutic intervention in high-burden settings.

## In Brief

Multi-omic profiling of acute illness in children across Africa and Asia reveals shared and syndrome-specific responses in immune, metabolic and lipid pathways, often shaped by malnutrition. These findings provide systems-level insights into paediatric illness biology and highlight candidate pathways for targeted interventions in high-burden settings.

## INTRODUCTION

Acute illness remains a leading driver of childhood mortality and morbidity globally, with the highest burden concentrated in low- and middle-income countries, particularly in sub-Saharan Africa and South Asia^1,2^. Children in these settings often present with overlapping clinical syndromes such as malnutrition, diarrhoea, malaria, pneumonia, sepsis and HIV infection that interact with underlying vulnerabilities in nutritional, immune and metabolic function^3–6^. Despite advances in clinical management, the biological mechanisms that shape acute illness and recovery remain poorly defined.

Acute illness triggers a cascade of systemic responses including immune activation, metabolic reprogramming and tissue remodeling. At the point of hospital admission, this is often reflected by elevated inflammatory and immune activation markers, and sometimes dysregulated immune function. For instance, children with septic shock exhibit sustained activation of the innate immune system and inflammation during hospitalisation, accompanied by suppression of adaptive immunity – patterns that are less pronounced in systemic inflammatory response syndrome (SIRS)^7^.

Metabolic alterations are also evident, for example, in Ugandan children with complicated severe malnutrition, baseline hypoleptinemia and decreased levels of high-molecular weight adiponectin were associated with inpatient mortality^8^. Similarly, Kenyan and Malawian children with complicated severe malnutrition who died during hospitalisation exhibited heightened systemic inflammation, reduced amino acids and long-chain lysophospholipids, and disturbances in mitochondrial metabolism^9^. In adults admitted with sepsis, greater metabolomic disturbance was associated with elevated mortality risk^10^. Infections can also amplify metabolic derangements in malnourished children. Kenyan and Ugandan children with HIV and concurrent severe wasting showed reduced zinc-alpha-2-glycoprotein, adiponectin and butyrylcholinesterase, alongside elevated triglycerides, ketones and acylcarnitines, compared to those with severe wasting alone^11,12^.

Malnutrition impairs host defenses through coordinated disruption of immune and metabolic pathways. Studies in children from sub-Saharan Africa and South Asia show impaired mucosal barrier integrity, thymic atrophy, altered T-cell function and dysregulated cytokine signaling, often coexisting with chronic immune activation and elevated immunoglobulin levels^13–15^. These immune alterations occur alongside suppressed metabolic capacity and endocrine dysregulation, reflecting an energy-conserving immunometabolic state that undermines pathogen clearance and tissue repair^13,15^. While there is growing evidence of molecular mechanisms underpinning severe malnutrition, little is known about how biological responses vary across the nutritional gradient or differ by clinical syndrome. For instance, in Kenyan children presenting with severe malaria, malnutrition was associated with a four-fold increased odds of true severe disease, and HIV infection with nearly ten-fold increased odds, while both alongside invasive bacterial infection were independently associated with higher mortality risk, underscoring the potential modulatory impact of nutritional and infectious comorbidities on syndrome-specific outcomes^3^.

Further, in infectious diseases, widespread disruption of immune-related processes has been reported in children, notably early innate immune activation, preserved adaptive responses, and, in some cases, dysregulated inflammation manifesting as multisystem inflammatory syndrome^16–18^. Severe disease maybe characterised by hyperinflammation, endothelial activation and complement-driven vascular injury^19,20^, highlighting the interplay between immune and vascular dysfunction. These findings underscore the profound disturbances to systemic biology that result from acute illness, whether driven by infection, malnutrition or their combination. However, most existing studies have examined isolated biomarkers or single pathways, limiting our ability to understand how these biological responses interact across systems and syndromes. The extent to which anthropometric status modulates these interactions, and how they differ from biological profiles in well children at risk of illness, remains insufficiently understood. Moreover, data on paediatric acute illnesses and the underlying biological processes remain sparse and scanty, particularly in resource-constrained settings.

In contrast, the biological underpinnings of disease in high-income settings have been extensively mapped through large-scale, multi-omic initiatives. Projects such as the UK Biobank (https://www.ukbiobank.ac.uk/use-our-data/), GTEx^21^ and various disease-specific studies have used transcriptomic, proteomic^22,23^, metabolomic and genomic^10,24^ approaches to profile biological variation in adult populations, transforming understanding of disease trajectories, treatment responses and aging. These resources have shaped precision medicine and driven the development of comprehesive molecular reference maps for adult diseases. However, similar large-scale biological data is lacking for children in low- and middle-income countries, despite the disproportionate burden of acute illness and mortality in these populations. Thus, a comprehensive, system-wide characterisation of illness in high-burden paediatric populations is urgently needed to complement existing resources and address critical gaps in global health biology.

To address these gaps, we conducted a multi-omic analysis of children hospitalised with acute illness across multiple sites in sub-Saharan Africa and South Asia. By systematically profiling proteomic, metabolomic, lipidomic and haematologic data, we aimed to: (i) identify shared and syndrome-specific biological pathways associated with acute illness; (ii) evaluate how malnutrition modifies systemic responses within syndrome-associated processes; and (iii) compare biological systems in acutely ill children to those in well community peers to examine how these systems interact in the two physiological contexts. This systems-level framework provides new insight into the molecular pathophysiology of childhood illness and has the potential to better understand multi-morbidity in children to inform more effective therapeutic strategies in high-burden settings.

## RESULTS

### Study participant characteristics at hospital admission

Figure 1A shows the selection of study participants. The characteristics of study children (n=1008) are presented in Table 1 and stratified by enrolment anthropometry in Table S1. The median age was 10.6 months (IQR: 6.5, 15.8) and 57% were males. Diarrhoea, anaemia and pneumonia were the most common presentations, and frequently co-occurred with malnutrition (Table 1, Figure 1B).

**Figure 1.**
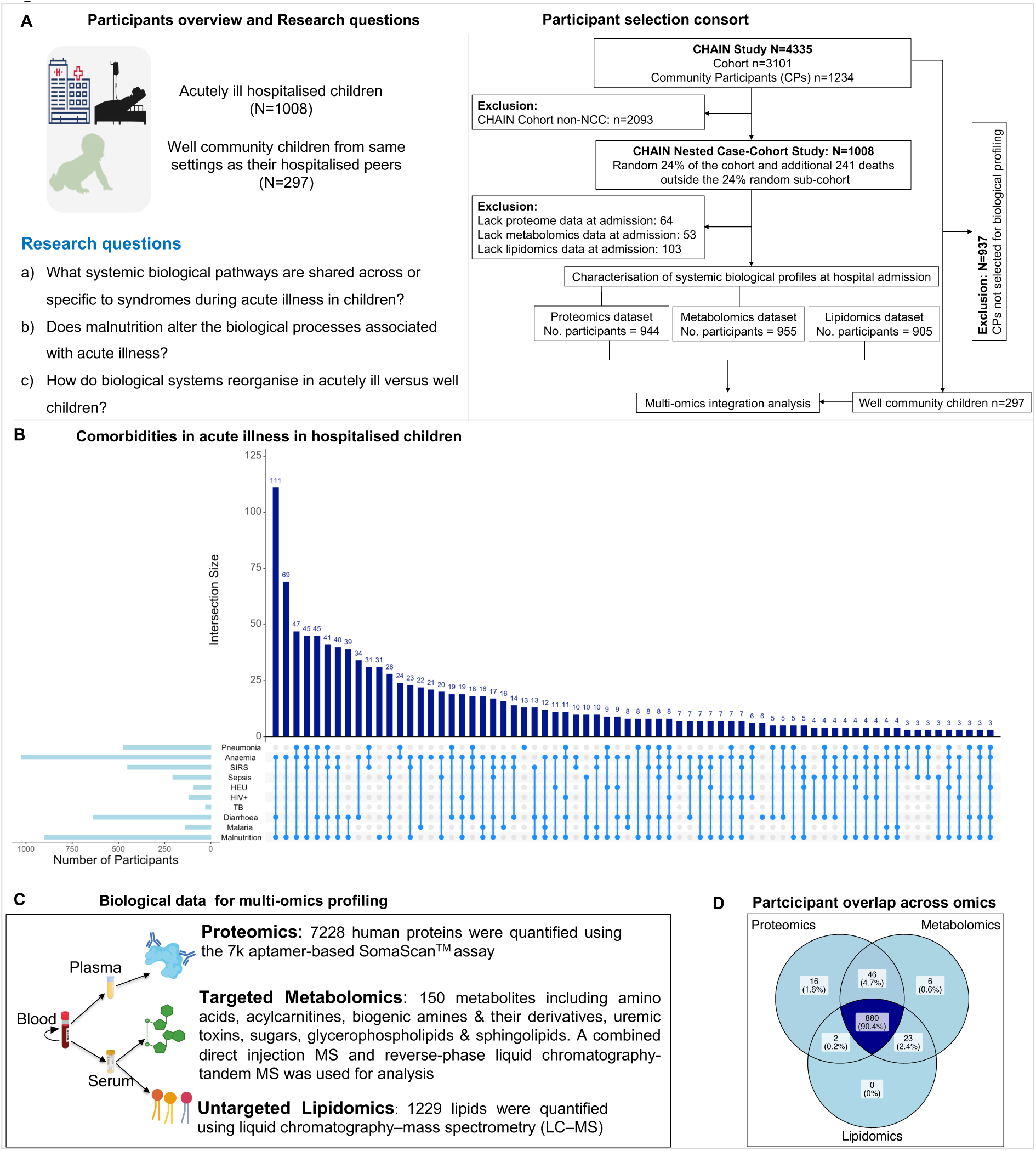
Study overview, design, participant consort and datasets. **(A)**. Study overview highlighting the number of acutely ill hospitalised and well community participants included in this study and the research questions addressed by the study. A total of 1008 acutely ill hospitalised children and 297 matched well community children were included in the current study. These children are drawn from 4 Sub-Sahara African and 2 South Asian countries. For characterisation of the baseline demographic, clinical, haematologic and biochemical features, all the 1008 hospitalised children were included. To understand the systemic biological profiles of acute illness in children, the study analysed 944, 955 and 905 children to unravel proteomic, metabolomic and lipidomic profiles respectively. Children who lacked these omic datasets were excluded from this analysis. **(B)**. Clinical comorbidities among acutely ill children at hospital admission. UpSet plot illustrating the frequency and patterns of clinical syndrome co-occurrences among children at hospital admission. The horizontal bars on the left represent the total number of participants diagnosed with each clinical syndrome. The dots connected by lines underneath show which syndromes are being combined. The vertical bars represent the size of participant intersections, showing how many children presented with specific combinations of syndromes. The most frequent comorbidity patterns included malnutrition combined with diarrhoea and anaemia (n=111), malnutrition with anaemia (n=69), malnutrition with anaemia and pneumonia (47 children) and two four-way combinations involving malnutrition, anaemia, SIRS and pneumonia (n=45), as well as malnutrition, diarrhoea, anaemia and pneumonia (n=45). The plot highlights the high burden of overlapping clinical comorbidities in this acutely ill paediatric population. **(C)**. Omic data analysed in this study. The study analysed proteomics (7,228 human proteins), metabolomics (150 metabolites including amino acids, acylcarnites, biogenic amines and their derivatives, uremic toxins, sugars, glycerophospholipids and sphingolipids) and lipidomics (lipid map categories analysed were glycerolipids, glycerophospholipids and sphingolipids) datasets. **(D)**. Participant overlap across proteomics, metabolomics and lipidomics datasets. 880 (90.4%) children had all the three omic datasets.

**Table 1.**
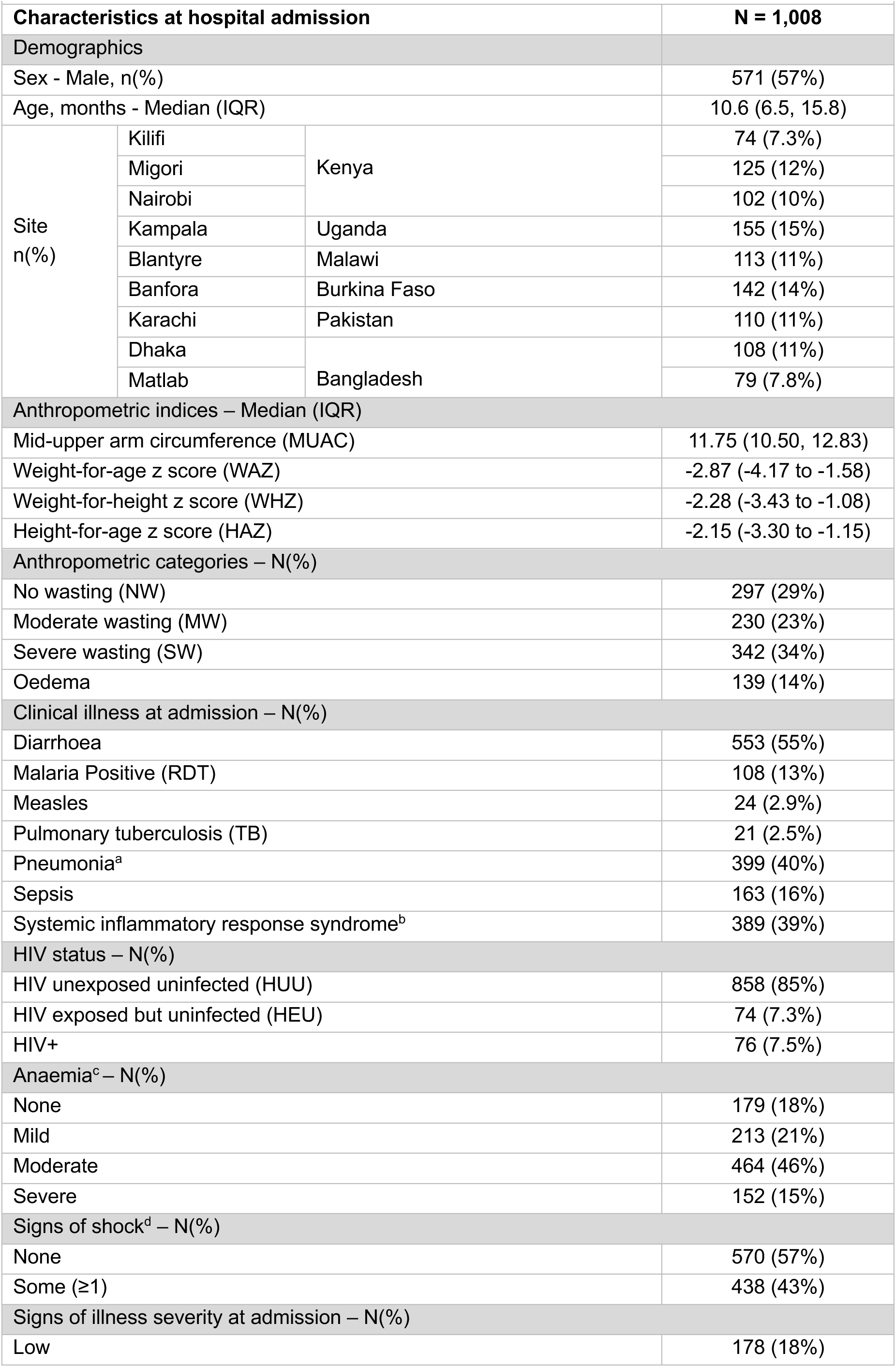
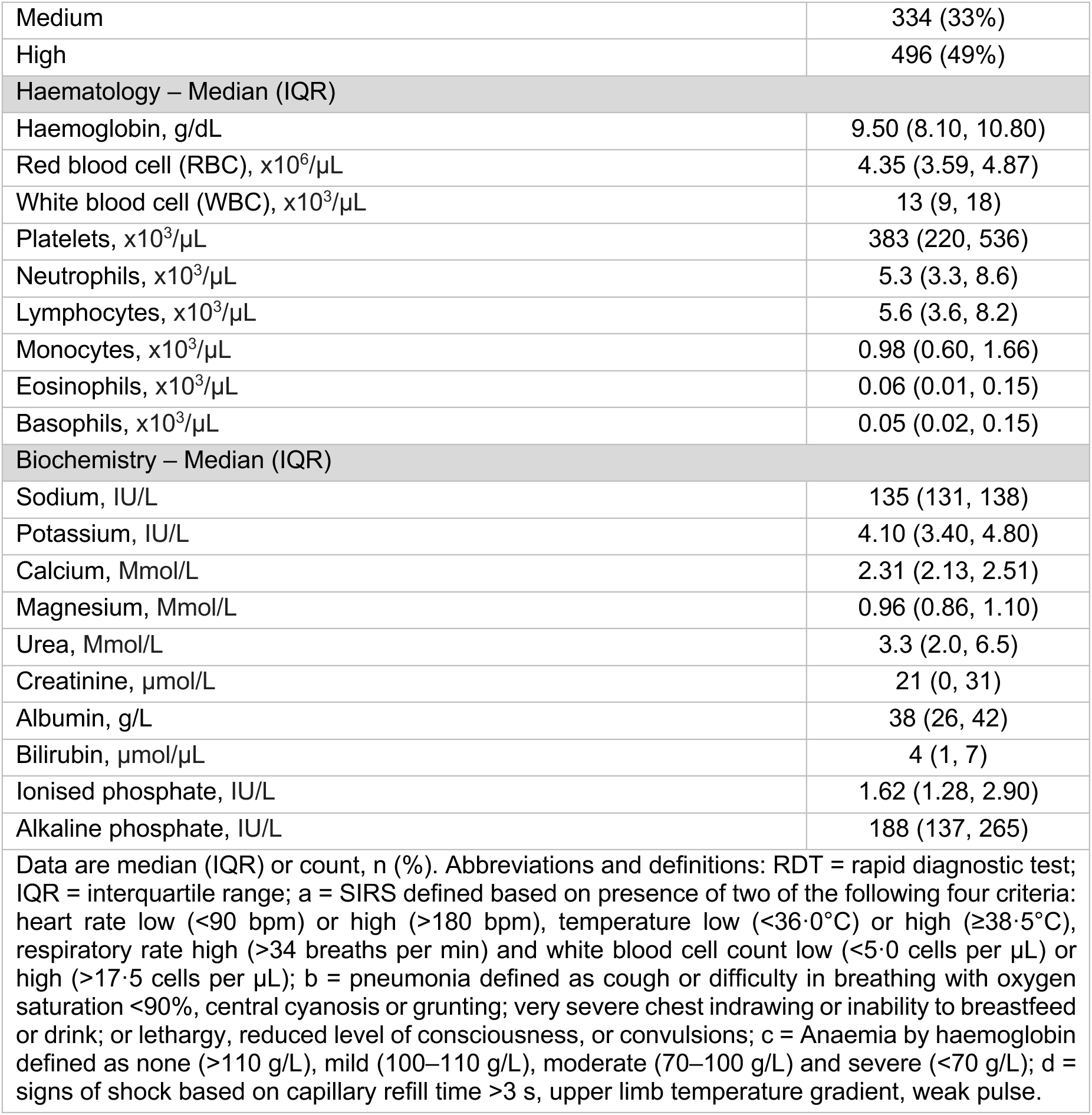
Study participants characteristics at hospital admission

| Characteristics at hospital admission |  |  | N = 1,008 |
| --- | --- | --- | --- |
| Demographics |  |  |  |
| Sex - Male, n(%) |  |  | 571 (57%) |
| Age, months - Median (IQR) |  |  | 10.6 (6.5, 15.8) |
| Site<br>n(%) | Kilifi | Kenya | 74 (7.3%) |
|  | Migori |  | 125 (12%) |
|  | Nairobi |  | 102 (10%) |
|  | Kampala | Uganda | 155 (15%) |
|  | Blantyre | Malawi | 113 (11%) |
|  | Banfora | Burkina Faso | 142 (14%) |
|  | Karachi | Pakistan | 110 (11%) |
|  | Dhaka | Bangladesh | 108 (11%) |
|  | Matlab |  | 79 (7.8%) |
| Anthropometric indices – Median (IQR) |  |  |  |
| Mid-upper arm circumference (MUAC) |  |  | 11.75 (10.50, 12.83) |
| Weight-for-age z score (WAZ) |  |  | -2.87 (-4.17 to -1.58) |
| Weight-for-height z score (WHZ) |  |  | -2.28 (-3.43 to -1.08) |
| Height-for-age z score (HAZ) |  |  | -2.15 (-3.30 to -1.15) |
| Anthropometric categories – N(%) |  |  |  |
| No wasting (NW) |  |  | 297 (29%) |
| Moderate wasting (MW) |  |  | 230 (23%) |
| Severe wasting (SW) |  |  | 342 (34%) |
| Oedema |  |  | 139 (14%) |
| Clinical illness at admission – N(%) |  |  |  |
| Diarrhoea |  |  | 553 (55%) |
| Malaria Positive (RDT) |  |  | 108 (13%) |
| Measles |  |  | 24 (2.9%) |
| Pulmonary tuberculosis (TB) |  |  | 21 (2.5%) |
| Pneumonia<sup>a</sup> |  |  | 399 (40%) |
| Sepsis |  |  | 163 (16%) |
| Systemic inflammatory response syndrome<sup>b</sup> |  |  | 389 (39%) |
| HIV status – N(%) |  |  |  |
| HIV unexposed uninfected (HUU) |  |  | 858 (85%) |
| HIV exposed but uninfected (HEU) |  |  | 74 (7.3%) |
| HIV+ |  |  | 76 (7.5%) |
| Anaemia<sup>c</sup> – N(%) |  |  |  |
| None |  |  | 179 (18%) |
| Mild |  |  | 213 (21%) |
| Moderate |  |  | 464 (46%) |
| Severe |  |  | 152 (15%) |
| Signs of shock<sup>d</sup> – N(%) |  |  |  |
| None |  |  | 570 (57%) |
| Some (≥1) |  |  | 438 (43%) |
| Signs of illness severity at admission – N(%) |  |  |  |
| Low |  |  | 178 (18%) |

|  |  |
| --- | --- |
| Medium | 334 (33%) |
| High | 496 (49%) |
| **Haematology – Median (IQR)** |  |
| Haemoglobin, g/dL | 9.50 (8.10, 10.80) |
| Red blood cell (RBC), $\times 10^6/\mu\text{L}$ | 4.35 (3.59, 4.87) |
| White blood cell (WBC), $\times 10^3/\mu\text{L}$ | 13 (9, 18) |
| Platelets, $\times 10^3/\mu\text{L}$ | 383 (220, 536) |
| Neutrophils, $\times 10^3/\mu\text{L}$ | 5.3 (3.3, 8.6) |
| Lymphocytes, $\times 10^3/\mu\text{L}$ | 5.6 (3.6, 8.2) |
| Monocytes, $\times 10^3/\mu\text{L}$ | 0.98 (0.60, 1.66) |
| Eosinophils, $\times 10^3/\mu\text{L}$ | 0.06 (0.01, 0.15) |
| Basophils, $\times 10^3/\mu\text{L}$ | 0.05 (0.02, 0.15) |
| **Biochemistry – Median (IQR)** |  |
| Sodium, IU/L | 135 (131, 138) |
| Potassium, IU/L | 4.10 (3.40, 4.80) |
| Calcium, Mmol/L | 2.31 (2.13, 2.51) |
| Magnesium, Mmol/L | 0.96 (0.86, 1.10) |
| Urea, Mmol/L | 3.3 (2.0, 6.5) |
| Creatinine, $\mu\text{mol/L}$ | 21 (0, 31) |
| Albumin, g/L | 38 (26, 42) |
| Bilirubin, $\mu\text{mol/L}$ | 4 (1, 7) |
| Ionised phosphate, IU/L | 1.62 (1.28, 2.90) |
| Alkaline phosphate, IU/L | 188 (137, 265) |
| Data are median (IQR) or count, n (%). Abbreviations and definitions: RDT = rapid diagnostic test; IQR = interquartile range; a = SIRS defined based on presence of two of the following four criteria: heart rate low (<90 bpm) or high (>180 bpm), temperature low (<36.0°C) or high ($\geq 38.5^\circ\text{C}$), respiratory rate high (>34 breaths per min) and white blood cell count low (<5.0 cells per $\mu\text{L}$) or high (>17.5 cells per $\mu\text{L}$); b = pneumonia defined as cough or difficulty in breathing with oxygen saturation <90%, central cyanosis or grunting; very severe chest indrawing or inability to breastfeed or drink; or lethargy, reduced level of consciousness, or convulsions; c = Anaemia by haemoglobin defined as none (>110 g/L), mild (100–110 g/L), moderate (70–100 g/L) and severe (<70 g/L); d = signs of shock based on capillary refill time >3 s, upper limb temperature gradient, weak pulse. | |

### Common and syndrome-specific biological processes characterise childhood response to acute illness

We examined the systemic biological landscape of acute illness at hospital admission in children with proteomics (n=944), metabolomics (n=955) and lipidomics (n=905) data (Figure 1C–D). Notably, 90% (n=880) of the children had all three omic datasets and characteristics of children with omics data were comparable to those with missing data (Table S2). Network analysis identified 45 protein modules (PM), 14 metabolite modules (MM), and 31 lipid modules (LM), reflecting distinct biological pathways (Figure 2A-C, Figure S1A-C,Tables S3– 5; see STAR methods). Protein modules ranged between 10 and 868 proteins, while metabolite and lipid ranged from 3–18 and 5–90, respectively (Figure S1A–C). Several modules were interconnected and formed supercusters (Figure S2A–B). The clinical syndromes examined in this study included malaria, diarrhoea, SIRS, TB, HIV, sepsis, anaemia and pneumonia, as defined in the STAR methods. We also provide a Shiny app via https://mudiboevans.shinyapps.io/Biological-Landscape-Multiomics/ to further facilitate in-depth exploration of distribution of omic modules and individual proteins, metabolites and lipids by demographics and across clinical syndromes analysed in the study.

**Figure 2.**
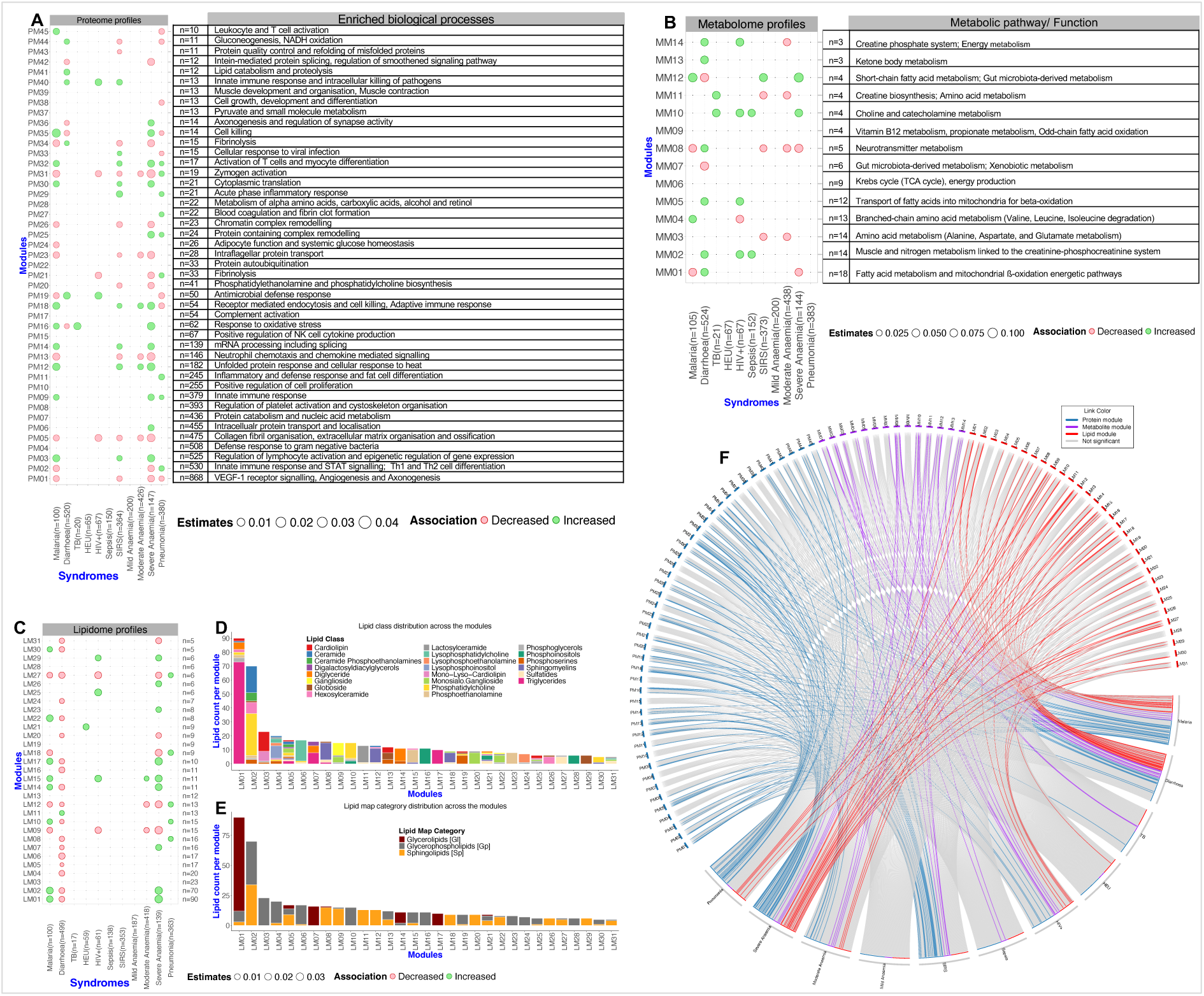
Associations between systemic biological modules and clinical syndromes at hospital admission. **(A–C)**. Bubble plots showing associations between clinical syndromes and omic modules. X-axes represent clinical syndromes, with *n* (in parentheses) indicating the number of children meeting the clinical definition for each syndrome. Y-axes represent modules from the proteome, metabolome and lipidome profiles respectively. Each bubble reflects the coefficient estimate from regression models, with size indicating the magnitude of the association. Red and green bubbles represent statistically significant negative and positive associations, respectively (p < 0.05). For all the regression models, the comparator group comprised children without the respective clinical syndrome. All regression analyses were adjusted for multiple comparisons using both Bonferroni and FDR corrections, as detailed in the STAR Methods. **(D–E)**. Lipid diversity across different modules. **(D)**. Stacked bar plots showing the distribution of lipids classes across the lipid modules. Each colour represents unique lipid species. **(E)**. Stacked bar plots showing the distribution of lipid categories across the lipid modules. Each colour represents unique lipid category. X-axes represent lipid modules while Y-axes represent lipid counts in a given lipid species constituting a specific category per module. **(F)**. Circular chord diagram summarising significant and non-significant associations between omic modules and clinical syndromes. Modules (proteomic, metabolomic and lipidomic) are displayed around the circumference and colored by omic layer. Curved edges denote associations where the colored lines represent statistically significant associations (p < 0.05; corrected for multiple testing as higlighted above) and grey lines indicate non-significant associations. Abbreviations: PM = protein module denoted by blue colour; MM = metabolite module represented by purple colour; LM = lipid module denoted by red colour; TB = tuberculosis; HEU = HIV-exposed uninfected; SIRS = systemic inflammatory response syndrome.

#### Proteomic responses

Despite clinical heterogeneity, common biological processes were observed across multiple syndromes. Many syndromes were marked by upregulation of processes related to inflammation, innate immunity and cellular stress (Figure 2A, F; Table S3). For example, modules involved in acute phase response, lymphocyte activation, unfolded protein response, and protein translation (e.g., PMs 3, 9, 12, 29, 30, 32, 40) were elevated across multiple syndromes (Figure 2A). Concurently, downregulation of biological processes essential for tissue repair and growth including angiogenesis, extracellular matrix organisation, ossification, chromatin remodelling, intraflagellar protein transport, phospholipid biosynthesis and post-translational modifications (PMs 1, 5, 20, 23, 26, 42) was observed across syndromes. These findings point to a systemic prioritisation of immune defense and stress adaptation over regeneration and growth-related processes during illness in children. The simultaneous suppression of neutrophil chemotaxis and chemokine-mediated signaling across multiple syndromes (PM13, Figure 2A) further suggests potential impairments in immune cell recruitment, highlighting a paradox in which heightened inflammatory activity may coexist with reduced cellular chemotaxis. This integrated response may reflect an adaptive but costly trade-off, limiting recovery and contributing to poor outcomes in paediatric acute illness.

In addition to these shared responses, we also observed syndrome-specific signatures (Figure 2A). Severe anaemia was associated with increased protein transport (PM06), pneumonia with elevated inflammatory and coagulation-related modules (PMs 11, 27), and diarrhoea with activation of lipid catabolic and proteolysis pathways (PM41). Syndrome-specific downregulated processes included glucose metabolism (PM24) in malaria; cell growth (PM38) in pneumonia and quality control of synthesised proteins (PM43) in SIRS.

Several modules exhibited divergent association patterns depending on the syndromes. For instance, PMs linked to oxidative stress response, T cell activation, receptor-mediated endocytosis and cell killing (PMs 16, 18, 35 and 45) were upregulated in malaria, TB and severe anaemia but downregulated in diarrhoea and pneumonia while glucose metabolism pathways, reflected in PM44 (gluconeogenesis and NADH oxidation) were elevated in diarrhoea but suppressed in SIRS and pneumonia. Fibrinolytic processes (PM21, PM34) were reduced in malaria, HIV, SIRS and severe anaemia but elevated in pneumonia and diarrhoea. Similarly, PM36 (axonogenesis) was selectively upregulated in severe anaemia and downregulated in diarrhoea. PM19 containing proteins indicative of antimicrobial defense response was upregulated in diarrhoea and HIV but decreased in malaria and pneumonia.

Moreover, processes linked to innate immune response (PM02) and proenzyme activation (PM31) were elevated in pneumonia but suppressed in malaria, HIV, SIRS, moderate and severe anaemia. Lastly, PM33, which includes proteins involved in antiviral response was increased in SIRS but reduced in pneumonia. Collectively, these divergent patterns of associations suggest context-specific immune and metabolic responses.

#### Metabolomic responses

Metabolomic analysis revealed both shared and syndrome-specific associations (Figure 2B, F, Table S4). Common metabolic associations included increased nitrogen and creatine metabolism (MM02), mitochondrial fatty acids oxidation (MM05) and catecholamine metabolism (MM10), reflecting increased bioenergetic demands. Across syndromes we also observed downregulation of specific amino acids including alanine, glutamine, methionine and hydroxy-lysine (MM03). In contrast, diarrhoea was uniquely marked by elevated ketone body metabolism (MM13) and suppression of gut microbiota-derived metabolism (MM07), suggesting altered gut microbial activity. Short-chain fatty acid metabolism (MM12) was decreased in diarrhoea but elevated in malaria, SIRS and severe anaemia. Fatty acid metabolism (MM01) was reduced in malaria and severe anaemia but upregulated in diarrhoea. Lastly, neurotransmitter and branched-chain amino acid metabolism (MM04 and MM08) and breakdown of amino acids to support the regeneration of ATP through the phosphocreatine system (MM11, MM14) varied by syndrome, highlighting widespread metabolic stress and compensatory adaptations (Figure 2B).

#### Lipidomic responses

Lipidomic network clustering demonstrated lipid class diversity within modules and distinct association patterns across syndromes (Figure 2C–F, Table S5). Particularly, in diarrhoea, which exhibited widespread lipid depletion across 18 modules (Figure 2G–K). This included suppression of triglycerides, phosphatidylcholines, sphingomyelins and gangliosides which are essential for energy storage and membrane integrity, indicating possible impaired dietary intake or lipid malabsorption. In contrast, pneumonia was predominantly associated with upregulation of lipid profiles including immune and inflammatory related lipid species like sphingomyelins, sulfatides and phosphatidylcholines (LM08, LM10, LM18, LM27). Malaria and severe anaemia showed increase in energy dense lipids species such as triglycerides and diglycerides (e.g., LM01, LM07, LM14, LM17). Across syndromes, we also noted consistent upregulation of glycerophospholipids (phosphoethanolamines [C32–C40], LM15) – key structural components of mitochondrial membranes, and sphingolipids (globosides [GB3: C34–C42], LM29) – enriched in lipid rafts and immune cell membranes. These alterations suggest membrane remodeling, immune cell activation and metabolic adaptation to stress during acute illness. Moreover, their concurrent elevation may reflect a shift towards catabolic metabolism. On the other hand, syndrome-specific associations included: LM21 (sulfatides, sphingomyelins, phosphatidylinositols, mono-lyso-cardiolipins; HEU), LM25 (phosphoglycerols, monosialo-ganglioside, mono-lyso-cardiolipins, cardiolipins; HIV), LM26 (hexosylceramides; severe anaemia), LM23 (phosphoethanolamines; severe anaemia), and LM11 (lactosylceramides; diarrhoea), suggesting context-specific lipidomic changes during illness.

Jointly, our omics analyses reveal that despite heterogeneous clinical presentations, many syndromes converge on shared systemic biological responses during acute illness. These include coordinated antimicrobial responses, protein synthesis, and fatty acid and amino acid catabolism to support energy mobilisation. Concurrently, multiple syndromes exhibited downregulation of processes critical for tissue maintenance and growth, such as extracellular matrix organisation, ossification, lipid and glucose metabolism and angiogenesis, and chemokine mediated signalling. While many associations were common across syndromes, some exhibited syndrome-specific signatures. For instance, diarrhoea was marked by global suppression of lipid metabolism and elevated lipid catabolic processes and proteolysis underscoring profound metabolic vulnerability in diarrhoea, while severe anaemia exhibited upregulation of protein transport and hexosylceramides. SIRS was characterised by reduced activity in protein quality control pathways. Distinct lipidomic profiles were also noted, with HIV and HEU showing unique enrichment of complex lipid species including cardiolipins, sulfatides and gangliosides, highlighting immunometabolic alterations unique to viral exposure, infection or antiretroviral therapy. Collectively, these findings delineate both convergent and syndrome-specific biological responses underlying paediatric acute illness.

### Malnutrition modifies syndrome-biological pathway associations

Having established that different clinical syndromes are associated with both shared and distinct biological pathways, we next examined whether these relationships were influenced by malnutrition status, defined by wasting (i.e., loss of muscle mass). Specifically, we examined whether the systemic biological processes linked to clinical syndromes differ between wasted and non-wasted children by assessing both the independent effect of wasting and its potential to modify these relationships.

After accounting for wasting (defined using mid-upper arm circumference, MUAC) associations between most clinical syndromes and biological modules remained robust (Figure 3A–I), suggesting that these biological processes are consistently associated with the respective syndromes regardless of a child’s level of wasting. However, for certain syndromes, specifically HIV some associations were no longer significant after adjusting for MUAC. Among children with HIV, the associations with PM19 (bacterial defense), PM21 (fibrinolysis), PM31 (zymogen activation) and MM04 (branched-chain amino acids metabolism) were no longer statistically significant after adjusting for MUAC (Figure 3D). These findings suggest that the observed associations were confounded by wasting and that HIV may not independently drive the respective biological pathways.

**Figure 3.**
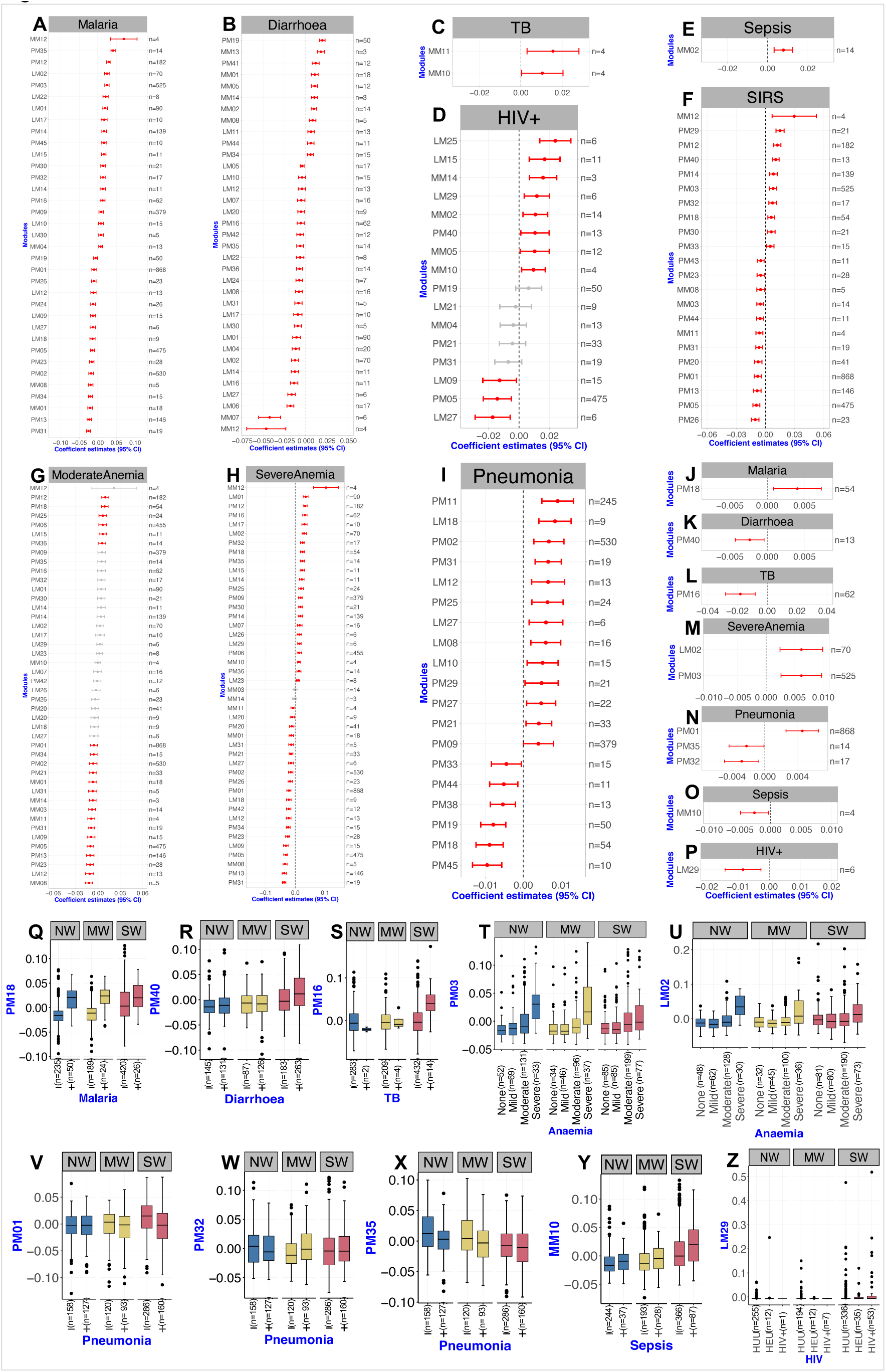
Wasting modifies certain biological pathway-syndrome associations. **(A–I)**. Forest plots showing β-coefficient estimates from additive models assessing associations between clinical syndromes and biological modules, adjusted for mid-upper arm circumference (MUAC). Each point (centre of the bars) indicate beta coefficient estimates for every unit increase in module concentration while error bars indicate 95% confidence interval. The red colour indicates significantly associated modules after MUAC adjustment. The number of features (proteins, metabolites or lipids) comprising each module is indicated on the right-hand side of each forest plot. **(J–P)**. Forest plots displaying β-coefficient estimates from interaction models testing the modifying effect of wasting (MUAC × clinical syndrome) on module expression. Results from additive and interaction models were corrected for multiple testing using Bonferroni and FDR methods, see the STAR Methods. **(Q–Z)**. Boxplots illustrating expression profiles of modules with significant interaction effects, stratified by syndrome status and anthropometric categories: NW (no wasting), MW (moderate wasting) and SW (severe wasting). Each box represents the interquartile range (IQR; 25th to 75th percentile), with the median indicated by the centre line. Whiskers extend to 1.5× the IQR and individual points beyond this range represent outliers. Abbreviations: MUAC = mid-upper arm circumference; PM = protein module; MM = metabolite module; LM = lipid module; TB = tuberculosis; HUU = HIV-unexposed uninfected; HEU = HIV-exposed uninfected; SIRS = systemic inflammatory response syndrome.

We found that malnutrition status modified several relationships between clinical syndromes and biological pathways. Specifically, interaction analysis revealed that the associations between some clinical syndromes and biological processes were dependent on childhood wasting (Figure 3J–P). For example, in malaria, PM18 linked to endocytosis and cell killing was elevated in wasted children without malaria, but remained consistently high across all anthropometric strata among those with malaria (Figure 3Q), suggesting a possible effect of wasting in the absence of malaria, which may be obscured by a dominant malaria-driven activation. In diarrhoea, expression of PM40 (antimicrobial response) was highest in severely wasted children suggesting synergistic upregulation (Figure 3R). We also noted that, while limited in sample size, severely wasted children with TB exhibited increased response to oxidative stress (PM16; Figure 3S).

In severe anaemia, wasting was associated with reduced expression of PM03 (lymphocyte activation; Figure 3M, T) and LM02 (lipid mobilisation; Figure 3M, U). In pneumonia, wasting attenuated pathways related to angiogenesis (PM01), T-cell activation (PM32), and cytotoxicity (PM35) suggesting impaired vascular and immune responses (Figure 3N, 3V-X). Sepsis-associated catecholamine-stimulated lipolysis (MM10) was also enhanced in wasted children (Figure 3O, Y). In HIV, wasting modified glycosphingolipid metabolism (LM29) with severely wasted children exhibiting increased globoside levels (Figure 3P, Z).

In summary, our results revealed that wasting modifies the relationships between systemic biological processes and specific clinical syndromes, and demonstrates that some pathways are prioritised over others during complicated severe wasting. Treatment strategies should therefore consider malnutrition status for optimal efficacy.

### Illness-specific and conserved co-regulation patterns in children

After examining how wasting influences the associations between clinical syndromes and biological pathways, we subsequently investigated how acute illness affects the co-expression patterns of individual proteins, metabolites and lipids. We compared hospitalised and well children using module preservation analysis (see STAR Methods). Demographics and anthropometric characteristics of community children are detailed in Table S6, with community children exhibiting better anthropometric indices than hospitalised peers. Among the 45 PMs in hospitalised children, 15 showed strong preservation (Z-summary >10) including PMs 1 to 11, 13, 14, 17 and PM30 (Figure 4A-B). Twenty-nine modules demonstrated moderate preservation (Z-summary; 2 to 10), while PM40 enriched for innate immune response and intracellular pathogen killing was not preserved (Z-summary <2). Strong preservations suggested that interactions among individual proteins and associated biological processes are conserved irrespective of illness status while moderate preservation reflected partial rewiring of these protein networks during acute illness. The lack of preservation for PM40 suggested active co-expression of specific proteins related to antimicrobial response during acute illness.

**Figure 4.**
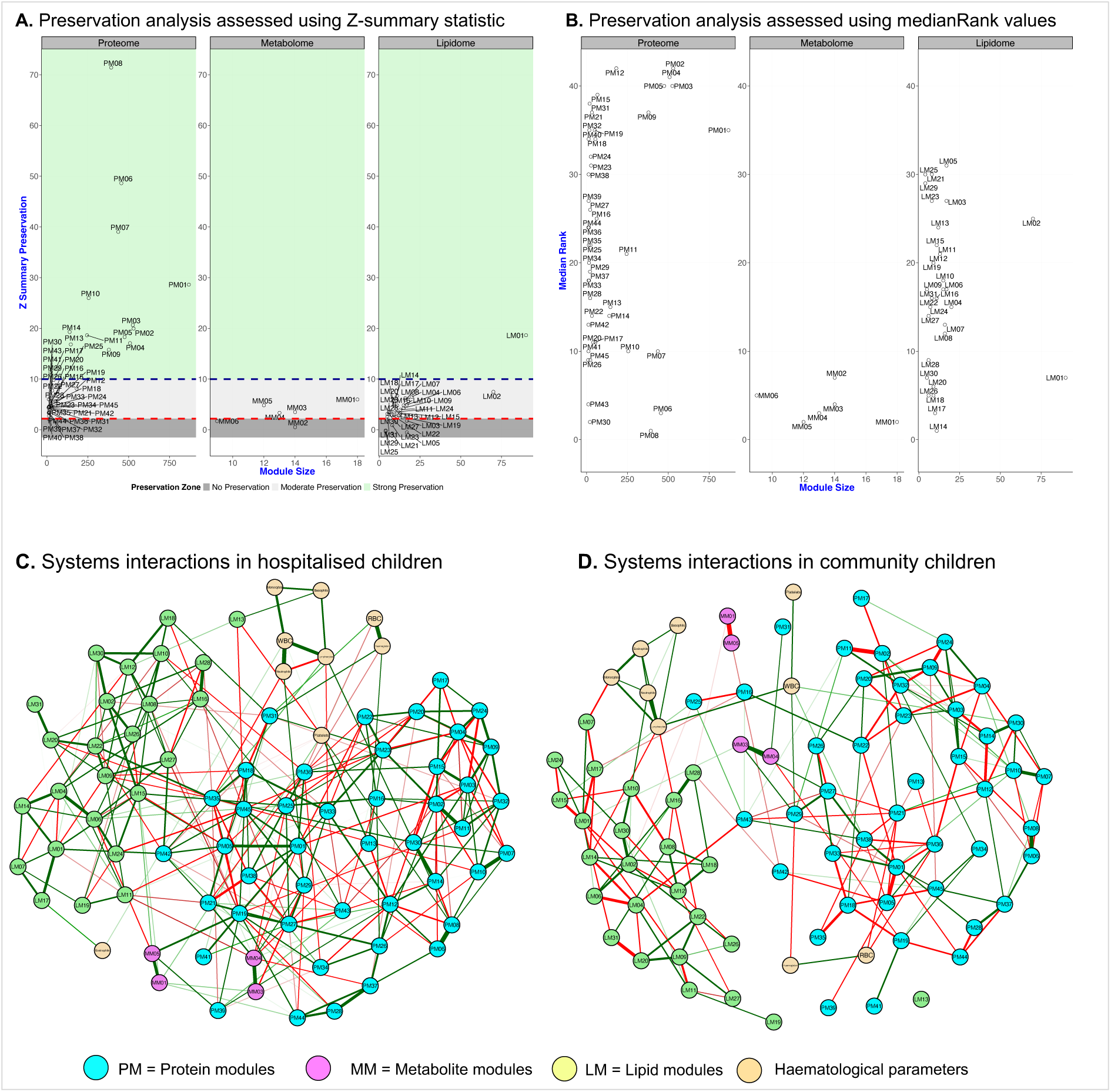
Conserved and acute-illnes induced co-expression patterns of biological features and systemic multi-omic network interaction across preserved modules. **(A)**. Module preservation assessed using the Z-summary statistic from WGCNA to evaluate whether the network structure of protein, metabolite and lipid modules derived from hospitalised children is retained in well community peers. The Y-axis show the Z-summary score, while the X-axis represent module size (i.e., number of features per module). Horizontal dashed lines denote standard interpretation thresholds: Z-summary >10 (green coloured zone) indicates strong preservation (preserved modules); 2 to 10 (light grey zone) suggests moderate preservation and <2 (dark grey zone – below the red line) indicates no preservation. Each bubble represents a module (e.g., PM01, MM01, LM01). The modules within green coloured regions exhibited strong preservation. These plots highlight that while a subset such as PM08 (proteins) and LM01 (lipids) showed strong preservation, most modules exhibited moderate or limited preservation, indicating context-dependent modulation of systemic biological networks during acute illness. **(B)**. To complement these findings, we also applied the medianRank statistic which is less influenced by module size to assess module stability. Lower medianRank values denote greater preservation across datasets. **(C–D)**. Systemic multi-omic network interaction for acutely ill hospitalised **(C)** and well community **(D)** children. The networks comprise only preserved biological modules, i.e., modules that strongly or moderately existed both in the acutely ill and well community children. Each node represents a module: protein (PM; cyan), lipid (LM; green), metabolite (MM; purple) or a haematological parameter (beige). Edges represent regularised partial correlations between module eigenfeatures or haematological markers; this approach allow meaningful comparison of biological networks arising from varying sample size. Green edges indicate positive correlations; red edges indicate negative correlations. Edge inclusion threshold of 0.06 and tuning parameter of 0.25 were applied to optimise the trade-off between model sparsity and sensitivity. Edge thickness reflects the magnitude of association. The hospitalised children’s network **(C)** shows a densely interconnected structure with strong cross-omic and haematological integration. In contrast, the well community children’s network **(D)** is modestly connected, with relatively less cross-omic interactions, possibly reflecting homeostatic independence among biological systems.

Due to analytical constraints, only six metabolite modules containing ≥9 features were assessed. None showed strong preservation (Figure 4 A–B). MM01 and MM05 (acylcarnitine metabolism and mitochondrial β-oxidation), and MM03 and MM04 (amino acid metabolism) were moderately preserved, while MM02 and MM06 (amino acid catabolism and TCA cycle activity) were not preserved. These finding suggest that protein catabolism and metabolic stress are hallmarks of acute disease. This is consistent with elevated circulating urea and creatinine previously observed in children at high risk of mortality^25^.

Lipid modules exhibited the most heterogeneous preservation profiles (Figure 4 A–B). LM01, dominated by triglycerides and other energy-related lipids was strongly preserved, implying a core role in both homeostatic and disease states. Twenty-four modules including LMs 2, 4, 6 to 12 among others were moderately preserved, while LMs 3, 5, 21, 25 and 29 were not preserved in well community children, indicating lipidomic network reorganisation specific to acute illness (Figure 4A–B, Table S5).

### Acute illness drives network rewiring and enhanced cross-omic coordination

After characterising the behaviour of individual biological processes in both homeostatic and acute disease states, we subsequently compared how these processes interact at the systems level during illness versus health. To assess the reorganisation of biological systems in acute illness, we constructed partial correlation networks using preserved modules, integrating omics and haematological features, and used node-level centrality metrics to evaluate shifts in biological coordination (Figure 4C–D, STAR methods). Networks derived from hospitalised children exhibited denser cross-omic connectivity than those from well children (Figure 4C–D, S3A–B). Node strength centrality was significantly increased in the illness network (Figure S3C). Nodes with marked increase in strength included PM12 (unfolded protein response), PM19 (antimicrobial response), PM27 (coagulation), PM35 (cell killing), PM31 (zymogen activation), PM42 (intein splicing – a post-translational modification), LM27 (sulfatides) and white blood cells. In contrast, eosinophils, sphingolipids (LM31) and monocytes had greater strength centrality in well children, suggesting shift in immune and lipid regulatory architecture during illness.

Closeness centrality, the closeness of each node to others, increased uniformly in the network of hospitalised children (p=2.55 × 10⁻¹⁵) supporting tighter biological interactions during illness. The greatest increases in closeness were observed for PM31 (zymogen activation), PM35 (cell killing), LM13 (cardiolipins, globosides, phosphoserines), PM18 (receptor-mediated endocytosis and cell killing), MM05 (acylcarnitine metabolism) and PM45 (leukocyte and T-cell activation). Others including PM43 (protein quality), LM07 (diglycerides and triglycerides), LM31 (sphingolipids), eosinophils and platelets had modest gains (Figure S3D).

Betweenness centrality showed bidirectional changes, suggesting a redistribution of system-level control points. Modules such as PM43, LM04, neutrophils and PM21 showed higher betweenness in community children (Figure S3E), suggesting homeostatic control roles. In contrast, PMs 1, 3, 12, 23, 33 and 35, and LMs 15 and 22 were more prominent in illness, indicating reallocation of key communication hubs under inflammatory stress.

Directional inversions of correlations further supported disease-driven rewiring. For example, neutrophils and lymphocytes, positively correlated in community children, showed a strong negative correlation in the hospitalised children, a shift supported by elevated neutrophil-to-lymphocyte ratio in ill children (p < 4.3 × 10⁻^48^; Figure S4). Elevated neutrophil-to-lymphocyte ratio has been reported as a prognostic marker in COVID-19 patients^26^ and associated with prolonged hospital stay in adults with myocarditis^27^. Other reversals included MM01 and MM05 (acylcarnitines) as well as LM01 and LM17 (triglycerides) and LM15 (phosphoethanolamines) which were positively correlated in hospitalised children but negatively correlated in community, supporting altered fatty acid oxidation, lipid mobilisation and alternative energy sources during illness. Additionally, LM04 (lysophospholipids) and LM14 (unsaturated diglycerides), and PM01 (angiogenesis) and PM05 (ECM and ossification) among others exhibited reversed relationships, underscoring a systemic reprogramming of biological interaction. In community children, monocytes were negatively correlated with LM07 (energy metabolism) but showed no connections with other modules in hospitalised children.

Collectively, these results demonstrate that acute illness induces extensive reorganisation of biological systems, characterised by heightened cross-omic connectivity, shifts in centrality control and directional inversion of key molecular relationships. These changes reflect an integrated, dynamic systemic response and may offer insight into biomarkers or targets for improved intervention during critical illness in children.

## DISCUSSION

This study presents a comprehensive systemic-level analysis of acute childhood illness, integrating proteomic, metabolomic, lipidomic and haematological data, from over 900 children in sub-Saharan Africa (Kenya, Uganda, Malawi, Burkina Faso) and South Asia (Pakistan and Bangladesh). We identified a core set of shared biological pathways across multiple syndromes, including inflammation, immunity, cellular stress, tissue repair and energetic pathways as well as syndrome-specific signatures linked to diarrhoea, HIV and anaemia. Wasting modified several immunometabolic and tissue repair responses in a syndrome-dependent manner. Compared to well community children, hospitalised children demonstrated greater cross-omic network integration, reversal of module correlations and elevated centrality in key immune, metabolic and coagulation pathways suggesting systemic rewiring during critical illness.

In resource-constrained settings, children admitted with acute illness frequently present with overlapping syndromes such as malaria, pneumonia, diarrhoea, anaemia or sepsis^1,6^. Such overlap is further compounded by malnutrition, particularly wasting^2^, micronutrient deficiencies^28^ and enteropathy^29^. In our cohort, multiple syndromes frequently co-occurred, with malnutrition present in a substantial proportion of children. These patterns highlight the intertwined nature of infection and undernutrition, emphasize the need to account for both comorbidity and anthropometric status when interpreting host biological responses.

Acute illness triggers immune activation, metabolic reprogramming and tissue remodelling, aimed at pathogen clearance and recovery. In our data, broad upregulation of acute-phase, antimicrobial defense, receptor-mediated endocytosis and cellular stress pathways reflected hallmarks of systemic inflammation. These arise from host responses to pathogen- and damage-associated molecular patterns, which drive innate immune and stress-adaptive programs^30,31^. Paradoxically, neutrophil chemotaxis and chemokine-mediated signalling were consistently downregulated, potentially indicating impaired immune cell recruitment or compensatory immunosuppression. Such decoupling of chemotactic signaling from systemic inflammation may reflect a reprogramming of host defense that prioritises containment and damage control over leukocyte trafficking, a phenomenon described in late-stage sepsis^32,33^ and severe malaria^34,35^. A recent study in Zimbabwean children with severe malnutrition demonstrated a shift in innate immune cell function, favouring bacterial containment over pro-inflammatory cytokine release^36^. Such adaptation may confer short-term survival benefits by limiting excessive inflammation at the expense of robust inflammatory signalling for pathogen clearance and development of immune memory. Concurrent downregulation of extracellular matrix organisation, ossification, and angiogenesis suggests compromised tissue repair and vascular integrity, consistent with a recent reports of microvascular damage and impaired growth recovery in severe illness^37,5^.

Acute illness imposes substantial energy demands due to immune activation, tissue repair and systemic stress. Fever alone can raise basal metabolic rate by ∼13% per °C, and bacterial endotoxins like lipopolysaccharide can elevate energy expenditure by up to 20%^38^. Meeting these demands requires metabolic flexibility and coordinated activation of energy-generating pathways^39^. Our findings reveal increased mobilisation of triglycerides and diglycerides, key energy storage lipids, in most syndromes but depletion in diarrhoea, likely reflecting impaired lipid absorption or enhanced catabolism. Upregulation of gluconeogenesis and accumulation of short-/medium-chain acylcarnitines indicated increased reliance on amino acid and fatty acid substrates, suggesting an adaptive shift in central metabolism during acute illness. The concurrent reduction in circulating amino acids across multiple syndromes further support the notion of heightened substrate catabolism to meet systemic energy needs. Additionally, elevations in ketone body metabolism, catecholamine-stimulated lipolysis and the accumulation of short-/medium-chain acylcarnitines, broadly suggest a shift toward lipid-derived energy sources, potentially compensating for insulin resistance or glucose scarcity during critical illness^40^. We also observed elevated short-chain fatty acids and gut microbiota-associated metabolites across multiple syndromes, which may reflect alterations in gut microbial activity. Given that short-chain fatty acids are primarily microbial fermentation products, their accumulation may signal alterations in microbiota composition or function during acute illness, potentially contributing to the broader metabolic changes observed in hospitalised children. However, their decreased detection in diarrhoea may point to gut microbiota loss or metabolic exhaustion.

We further noted reductions in long-chain acylcarnitines in multiple syndromes, reflecting incomplete β-oxidation or mitochondrial inefficiency. These findings, coupled with the consistent upregulation of oxidative stress response pathways, point to increased mitochondrial reactive oxygen species production and compensatory antioxidant responses. Mitochondrial dysfunction has been implicated in poor outcomes in both sepsis and severe malnutrition, as demonstrated in our recent study where inpatient mortality among children treated for complicated severe malnutrition was linked to systemic inflammation and profound metabolic disturbances^9^. Notably, in the current study, the differential levels of long-chain and short-/medium-chain acylcarnitine modules in well community versus hospitalised children indicates mitochondrial metabolic adaptations during acute illness. In hospitalised children, the positive correlation between long-chain and short-/medium-chain acylcarnitines suggests a coordinated accumulation of acylcarnitines across chain lengths, likely reflecting impaired β-oxidation, mitochondrial overload or inhibition of downstream acylcarnitine metabolism. Such accumulation patterns are increasingly recognised as markers of incomplete fatty acid oxidation during metabolic stress and have been associated with mitochondrial dysfunction in critically ill patients^10,41^. Mitochondrial dysfunction during acute illness may arise from multiple mechanisms, including increased energy demand, oxidative stress, substrate limitation and cytokine-mediated inhibition of mitochondrial enzymatic pathways. Pro-inflammatory cytokines (e.g., TNF-α and IL-6) can suppress mitochondrial biogenesis and impair β-oxidation capacity^42^, leading to the accumulation of partially oxidised fatty acid intermediates like acylcarnitines. Additionally, nutrient imbalances common during critical illness, such as carnitine depletion or coenzyme A insufficiency^43^, may further exacerbate acylcarnitine buildup. Conversely, in community children, the negative correlation between long-chain and short-/medium-chain acylcarnitines likely reflects efficient regulation of fatty acid flux, with dynamic partitioning and utilisation of short-, medium-, and long-chain fatty acids in a manner that maintains metabolic homeostasis. In well states, acylcarnitine levels are tightly controlled through flexible mitochondrial substrate switching and coordinated β-oxidation^44^, preventing accumulation of potentially toxic lipid intermediates. Together, these findings suggest that mitochondrial dysfunction is a central feature of systemic metabolic stress during acute illness in children, with altered acylcarnitine profiles serving as potential biomarkers of impaired mitochondrial energy metabolism.

Our findings demonstrated that malnutrition, specifically wasting, modifies host systemic response in a syndrome-specific manner consistent with prior studies demonstrating impact of malnutrition on infection^3^ and immune responses^4,14,45,46^. Severely wasted children with HIV had elevated gangliosides specifically globotriaosylceramides suggesting increased glycosphingolipid metabolism under combined metabolic and infectious stress. Dyslipidemia has been described in children with HIV within sub-Saharan Africa^47–49^ and it is common in stunted children^50,51^ and those with low body mass index for age^52^. On the other hand, there was attenuated expression of lipid classes that support mitochondrial membrane structure, redox regulation and energy transduction in wasted children with severe anaemia. These implies that malnutrition impairs lipid mobilisation and mitochondrial adaptation consistent with prior reports of mitochondrial dysfunction in undernourished states^53^. Beyond lipid metabolism, wasting modified several key protein modules involved in angiogenesis, T cell activation, cytotoxicity, oxidative stress response and host defense. These effects were most pronounced in pneumonia, diarrhoea and TB, where wasted children displayed altered responses. For instance, in pneumonia, wasting was associated with attenuated angiogenesis and diminished expression of T cell activation and cytotoxic pathways and therefore may exacerbate immune suppression and hinder tissue regeneration during pulmonary infections. In contrast, wasted children with diarrhoea had heightened antimicrobial responses while those with sepsis had elevated catecholamine-stimulated lipolysis and fatty acid metabolism reflecting strained bioenergetics. Together, these findings underscore the compounded impact of wasting and infection on immune, metabolic and tissue repair responses during acute illness.

The rewiring of biological networks observed in hospitalised children compared to well peers suggests a systemic response to acute illness that involves heightened coordination and integration across molecular systems. Compared to well community children, hospitalised children demonstrated greater systemic coordination, tighter network integration and reversal of relationship structures among select module pairs indicating dynamic reorganisation of biological systems during critical illness. Specifically, the uniform increase in strength and closeness centrality within networks of hospitalised children reflects a globally more integrated biological architecture, consistent with previous reports of heightened cross-system coordination and network tightening in altered physiological states^54,55^. These shifts represent adaptive responses aimed at maintaining homeostasis under systemic stress but also signal vulnerability to decompensation when adaptive mechanisms are overwhelmed, potentially culminating in multisystem organ failure – a common trajectory in critically ill children in low-resource settings^56,57^.

This study represents one of the most comprehensive systems-level investigations of acute illness in children from resource-constrained settings, integrating multi-omic and haematological features across a broad spectrum of clinical comorbidities including malnutrition, malaria, diarrhoea, TB, HIV, sepsis, SIRS, anaemia and pneumonia. Leveraging a large, geographically diverse cohort from 9 sites across sub-Saharan Africa and South Asia, our analysis captures a wide range of socio-demographic and clinical variability, enhancing the generalisability of our findings. The inclusion of both acutely ill hospitalised children and well community-based peers from the same settings offers a rare comparative framework for distinguishing biological features of illness from background variation. Together, these data provide key insights into the molecular pathophysiology of acute childhood illness and support the development of biologically informed approaches for risk stratification, biomarker discovery and targeted therapeutic interventions.

### Limitations of the study

In this study, we acknowledge that clinical syndromes were defined using World Health Organisation criteria (see STAR Methods), which may lack disease specificity and contribute to overlapping comorbidities, a common phenomenon in paediatric population studies in high-burden settings. This may, in part, underlie the convergence of associations across multiple syndromes. Furthermore, analyses were restricted to a single time point (hospital admission), limiting insight into the temporal dynamics of biological responses during hospitalisation and recovery. While this study provides detailed insights into the biological landscape of acute illness at the time of hospital admission, it did not investigate the association between these biological processes and clinical outcomes such as in-hospital or post-discharge mortality. As such, the prognostic relevance of the observed systemic alterations remains to be explored in future outcome-focused analyses. Finally, although models were adjusted for key demographic variables (including age, sex and site), data on other potential confounders such as socioeconomic status and caregiver characteristics were not analysed, which may influence biological–clinical associations.

## RESOURCE AVAILABILITY

### Lead contact

Further information and requests for resources should be directed to and will be fulfilled by the lead contact, Evans O. Mudibo.

## Materials availability

This study did not generate new unique reagents.

## Data and code availability

Data supporting this study are available at Harvard Dataverse^58^ through this link: https://doi.org/10.7910/DVN/PGTZNO. Due to sensitive participant-level information, access is restricted and requires approval from the KEMRI-Wellcome Trust Research Programme Data Governance Committee. Instructions for data requests and access are provided on the Harvard Dataverse landing page via the above link.

Summary visualisations (e.g., histograms, boxplots and principal component analysis plots) of distribution of omic modules and individual proteins, metabolites and lipids are available at https://mudiboevans.shinyapps.io/Biological-Landscape-Multiomics/.

Analysis code used in this study is available at the Harvard Dataverse^58^: via https://doi.org/10.7910/DVN/PGTZNO. All tools used are listed in the Key Resources Table.

## ACKNOWLEDGMENTS

We thank the CHAIN Network for their invaluable contributions to this study, including provision of clinical and biological data. The primary CHAIN study was funded by the Bill & Melinda Gates Foundation [INV-000791 and OPP1131320 to J.A.B. and J.L.W.]. For the purpose of open access, the CHAIN Network has applied a CC BY public copyright license to any author-accepted manuscript version arising from this submission.

Additional support was provided by the National Institute for Health and Care Research [NIHR201813 to A.J.P., G.B.G., P.K., J.A.B., K.D.T., J.M.N., and B.O.S.] and the Medical Research Council Department for International Development-Wellcome Trust Joint Global Health Trials scheme [MR/M007367/1 to J.A.B.]. E.O.M. is a PhD candidate jointly affiliated with KEMRI-Wellcome Trust Research Programme and Wageningen University, funded by NIHR [NIHR201813]. A.J.P. is supported by Wellcome [108065/Z/15/Z] and J.M.N. holds a Wellcome Trust Intermediate Fellowship [222967/B/21/Z]. The funders had no role in study design, data collection and analysis, decision to publish, or manuscript preparation.

## AUTHOR CONTRIBUTIONS

Conceptualisation of the study: E.O.M., J.A.B., G.B.G., J.M.N.

Design of the nested case-cohort study within CHAIN: K.D.T., M.M.N., J.M.N. Data analysis and interpretation: E.O.M., J.A.B., G.B.G., J.M.N.

J.A.B., G.B.G., and J.M.N. supervised the analysis, guided interpretation, and critically reviewed the manuscript. E.O.M wrote the original draft.

Equal contribution: J.A.B., G.B.G., and J.M.N.

J.A.B. and J.L.W. were the principal investigators of the original CHAIN Network study.

C.S., A.J.P., K.D.T., C.L., B.J., A.K., R.H.J.B., C.J.M., and J.L.W. critically reviewed the manuscript.

All authors approved the final manuscript and had full responsibility for the decision to submit for publication.

## SUPPLEMENTAL INFORMATION

Document S1. Figures S1–S5 and Table S1–S6

Table S7. List of proteins, metabolites and lipids analysed in the study

Table S8. Intra-modular connectivity for proteins, metabolites and lipids networks

## STAR★METHODS

### EXPERIMENTAL MODEL AND STUDY PARTICIPANT DETAILS

#### Study design, setting and population

This study is a secondary analysis nested within the Childhood Acute Illness and Nutrition (CHAIN) Network, a prospective cohort study designed to investigate biomedical and social determinants of mortality in acutely ill young children^59^. Between November 20, 2016 and January 31, 2019, CHAIN enrolled children aged 2 to 23 months admitted to hospitals in sub-Saharan Africa (Kenya: Kilifi, Mbagathi, and Migori County hospitals; Uganda: Mulago National Referral Hospital; Malawi: Queen Elizabeth Hospital; and Burkina Faso: Banfora Referral Hospital) and South Asia (Pakistan: Civil Hospital; and Bangladesh: Dhaka and Matlab Hospitals).

Children were hospitalised with various acute conditions, including malaria, diarrhoea, tuberculosis (TB), sepsis, systemic inflammatory response syndrome (SIRS), anaemia and pneumonia. Enrolment was based on mid-upper arm circumference (MUAC) into three anthropometric groups; not wasted: MUAC ≥12.5 cm for age ≥6 months or MUAC ≥12.0 cm for age <6 months, moderately wasted: MUAC 11.5 cm to <12.5 cm for age ≥6 months or MUAC 11.0 cm to <12.0 cm for age <6 months, and severely wasted or kwashiorkor: MUAC <11.5 cm for age ≥6 months or MUAC <11.0 cm for age <6 months or bilateral pedal oedema unexplained by other medical causes. In total, 3101 hospitalised children were enrolled. Additionally, well children (n=1234) from the same communities as the hospitalised cohort were seen at a single time point to establish community norms. Standardised procedures across all sites ensured comprehensive and harmonised collection of clinical, demographic, anthropometric and biological data (blood, stool and rectal swabs)^59^. Samples were archived at the Kilifi biobank in Kenya at -80° C.

This analysis is nested within the CHAIN case-cohort (CHAIN NCC) multiomic mortality sub-study which included a random sub-cohort of 24% (n=767) of CHAIN participants, all deaths not in the sub-cohort (n=241) and randomly selected community participants (n=290)^60^. Therefore, CHAIN NCC is comprised of 1008 hospitalised children. Multi-omic data available in CHAIN NCC include proteomics, metabolomics, lipidomics, faecal microbiome profiles, enteric pathogens detection and biomarkers of enteropathy^60^. For this analysis, we focused on data measured at hospital admission and specifically the study analysed proteomics, metabolomics and lipidomics data. The study excluded children missing proteomic (n=64), metabolomic (n=53) and lipidomic (n=103) data, as shown in Figure 1A.

Sex in this study was defined based on the biological attribute at birth (male or female). While no sex stratified analyses were performed, all our statistical models were adjusted for sex to account for its influence on biological processes, as detailed in the statistical methods section.

#### Ethics

Ethical approvals were obtained from all participating site institutions and collaborators, as well as the University of Oxford. Written informed consent was obtained from caregivers for each child. The study protocol received ethical clearance from: Oxford Tropical Research Ethics Committee (UK); Kenya Medical Research Institute (Kenya); University of Washington and Oregon Health and Science University (USA); Makerere University School of Biomedical Sciences Research Ethics Committee and the Uganda National Council for Science and Technology (Uganda); Aga Khan University (Pakistan); International Centre for Diarrhoeal Disease Research, Bangladesh (icddr,b); The University of Malawi and COMREC, Kamuzu University of Health Sciences (Malawi); University of Ouagadougou and Centre Muraz (Burkina Faso); the Hospital for Sick Children (Canada); and the University of Amsterdam (The Netherlands).

## METHOD DETAILS

### Clinical syndromes and definitions

Syndromes were identified based on standard clinical diagnostic criteria:

**Malaria**: Positive CareStart HRP2/pLDH rapid diagnostic test.

**Diarrhoea**: Passage of ≥3 loose or watery stools in 24-hour^61^.

**Systemic Inflammatory Response Syndrome (SIRS)**: at least two of: heart rate (<90 or >180/min), temperature (<36^0^C or ≥38·5^0^C), respiratory rate (>34 breaths per minute) and white blood cells (WBC) (<5 or >17·5 x10^9^/l)^62^.

**Anaemia**: Categorised by WHO thresholds^63^: none (haemoglobin; Hb >11g/dl), mild (Hb ≥10 to 11g/dl), moderate (Hb ≥7 to <10g/dl) and severe (Hb <7g/dl).

**Pneumonia**: WHO 2013 criteria^61^, including cough or difficulty breathing plus one or more of: central cyanosis, oxygen saturation <90%, chest indrawing, inability to feed, persistent vomiting or altered consciousness.

**Tuberculosis (TB):** Receiving treatment for TB.

**Human Immunodeficiency Virus (HIV)**: Polymerase chain reaction (PCR) or antigen testing per national guidelines.

### Baseline characteristics of the study participants

Baseline characteristics of the study children at hospital admission, including demographic (age, sex, and site of enrolment), anthropometry, clinical syndromes, complete blood count and clinical biochemistry were summarised using median with interquartile ranges for continuous variables and proportions for categorical variables (Table 1). Characteristics were stratified by enrolment anthropometric categories (no-, moderate- and severe wasting; Table S1). We also profiled the multimorbidity status of acute illness in the study children using the UpSet R package v1.4.0^64^.

### Laboratory analyses and data preprocessing

The study used plasma proteomics and serum metabolomics and lipidomics data from the CHAIN NCC sub-study as previously described^60^. All quantified features are listed in Table S7.

#### Plasma Proteomics

Plasma human proteins (n=7,228) were quantified using the 7k aptamer-based SomaScan^TM^ assay v4.1 (SomaLogic, Inc. USA)^60,65^. Raw text-based ADAT files were processed with the readat R package^66^. Protein data were log-transformed, mean-centred and scaled to unit variance. No missing values were reported by the SomaScan^TM^ assay.

#### Targeted Serum Metabolomics

A custom assay combining direct injection mass spectrometry and reverse-phase liquid chromatography-tandem mass spectrometry quantified 150 serum metabolites at The Metabolomics Innovation Centre (TMIC), Canada^60^. Metabolites included amino acids, acylcarnitines, biogenic amines and their derivatives, uremic toxins, sugars, glycerophospholipids and sphingolipids. Metabolites below detection in ≤30% of samples (n=6) were imputed using the lowest concentration of the metabolite divided by the square root of two. Lipids already quantified using untargeted lipidomics (14 lysophosphatidylcholines, 10 sphingomyelins and 10 phosphatidylcholines) were excluded leaving 116 metabolites which were log-transformed, mean-centred and scaled to unit variance.

#### Untargeted Serum Lipidomics

Lipidomic profiling was performed using liquid chromatography–mass spectrometry (LC–MS) following protein precipitation-based extraction^60,67^. Of the 1,229 lipids quantified, 601 with zero values were excluded. The remaining 628 were log-transformed, mean-centred and scaled to unit variance.

Data distribution was visualised using histograms, boxplots and Principal Component Analysis (PCA) plots, stratified by sex, site, anthropometric status and clinical comorbidities. Visualisations are accessible via: https://mudiboevans.shinyapps.io/Biological-Landscape-Multiomics/.

### Covariates

Key covariates included age, sex, MUAC and study site. Site was included as a random effect in the statistical models. Participants were not stratified by race, ethnicity or other social groupings.

### Construction of the biological correlation networks

To construct molecular networks across proteomics, metabolomics and lipidomics data, we employed Weighted Gene Correlation Network Analysis (WGCNA)^68^, an unsupervised systems biology approach that identifies modules of highly correlated features while assuming an underlying scale-free network topology; a characteristic typical of biological systems^69^. A network is considered to approximate scale-free topology when the correlation coefficient (R^2^) exceeds 0.8. Prior to network construction, we conducted quality checks on both features and samples, which confirmed that the data met the requisite criteria for WGCNA analysis. Using the WGCNA R package^68^ (v1.73), we first determined the optimal soft-thresholding power (β) by evaluating a range of powers from 1 to 50. An optimal β in network construction helps to achieve network sparsity and scale-free topology. Among the range of potential β that achieved a scale-free topology fit (R² ≥ 0.80), we selected the optimal β for each omic dataset (proteomic, metabolomic and lipidomic data) by choosing the value that resulted in a network with the fewest unassigned features and minimum mean connectivity. For proteomic, metabolomic and lipidomic datasets we selected β of 12, 3 and 5 respectively. Following selection of the optimal β, pairwise correlation matrices were generated for each of the omic layer. For proteomics, we computed pairwise biweight mid-correlations *𝑠*_𝑖𝑗_ = bi𝑐𝑜𝑟(*𝑥*_𝑖_, *𝑥*_𝑗_) while for metabolomics and lipidomics we computed pairwise correlations based on the Pearson’s correlations, *𝑠*_𝑖𝑗_ = 𝑐𝑜𝑟(*𝑥*_𝑖_, *𝑥*_𝑗_) – this was informed by our optimisation procedure for selecting ideal β. These correlation matrices were then converted into an adjacency matrix through power transformation (𝑎ᵢⱼ = |𝑠ᵢⱼ|^β), which emphasises strong positive correlations while suppressing weaker or negative ones, enhancing the signal-to-noise ratio in network construction.

Networks were built using a signed approach to preserve the directionality of relationships – for better biological interpretation, and modules of tightly co-expressed features were identified using hierarchical clustering via the *blockwiseModules* function. The minimum module size was set to 10, 3 and 5 features for proteomics, metabolomics and lipidomics data respectively. The entire dataset for each omic layer was analysed as a single block (maxBlockSize = 20,000) and closely related modules were merged using a dissimilarity threshold of 0.25. This approach allowed us to infer biological pathways rather than focusing solely on individual biomarkers^70^, while also reducing high-dimensional omics data into biologically meaningful modules for downstream analysis. We have previously applied the WGCNA approach to the SomaScan® proteomic data to identify protein modules associated with clinical outcomes^5^.

### Biological functional annotation

Protein modules were functionally annotated using Gene Ontology (GO) enrichment using DAVID (v2024q4)^71^, WEB-based Gene SeT AnaLysis Toolkit (WebGestalt)^72^ 2024 version and STRING^73^ (v12.0) with *Homo sapiens* as the background. Significant enrichment was defined using Bonferroni (for DAVID) or FDR adjusted p-values < 0.05 (WebGestalt and STRING). Further, hub proteins, representing highest connectivity core drivers of module structure, were identified by intra-modular connectivity from WGCNA to better characterise biological mechanisms (Table S3, S8).

Metabolite and lipid modules were annotated based on known biological functions of their constituent molecules (Table S4 - S5).

### Statistical analysis

The analytical approaches employed in this study are detailed in the following section and the link to the accompanying codes is available in the Key Resources Table.

## QUANTIFICATION AND STATISTICAL ANALYSIS

### Association between clinical syndromes and omic modules

Each module was summarised using its eigenfeature, the first principal component representing the overall behaviour of the module, derived via singular value decomposition. Associations between module-eigens (ME) and clinical syndromes were assessed using inverse probability-weighted (IPW) linear mixed-effects models implemented in the lmerTest R package (v3.1-3)^74^ and adjusted for age, sex, illness severity and random effects by site (Equation 1). Clinical syndromes included: malaria, diarrhoea, TB, HIV, sepsis, SIRS, anaemia (mild, moderate, severe) and pneumonia. IPW helped account for potential selection bias within CHAIN NCC which was enriched for mortality relative to the parent cohort. Multiple testing was controlled using Bonferroni and FDR methods, and significance was set at p-value < 0.05.

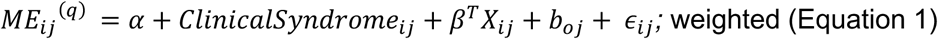

where 𝑀𝐸*_ij_*^(*q*)^ = module eigenfeature *q* of the respective omic layer; 𝐶𝑙𝑖𝑛𝑖𝑐𝑎𝑙𝑆𝑦𝑛𝑑𝑟𝑜𝑚𝑒*_ij_* = binary indicator for presence (1) or absence (0) of a specific clinical syndrome; 𝑋*_ij_* = vector of covariates (age, sex, illness severity); *β* = corresponding vector of coefficients for covariates; 𝑏*_oj_* = is a random intercept for site; 𝜖*_ij_* = is the residual error term.

### Influence of malnutrition on syndrome-associated modules

Following the identification of modules associated with clinical syndromes, we subsequently examined whether wasting (defined by MUAC) modified these relationships. To examine the effect of wasting on syndrome-module associations, we used additive and interaction models with MUAC as a continuous variable. Additive models (Equation 2) tested the confounding effect of wasting on these relationships while interaction models (Equation 3) evaluated the modification effect of wasting, with log-likelihood tests specifying which models had better fit. A significant likelihood ratio test supported the interaction models as the more appropriate. Models were adjusted for covariates (including age, sex, illness severity, oedema and site included as random effect), incorporated IPW, and applied Bonferroni and FDR corrections to control for multiple testing, with statistical significance set at p < 0.05.

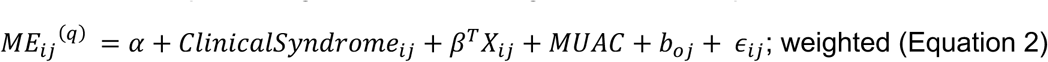

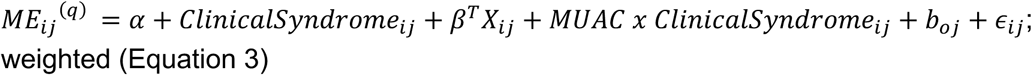

where 𝑀𝐸*_ij_* = module eigenfeature *q* of a given omic layer associated with a particular syndrome from Equation 1; 𝐶𝑙𝑖𝑛𝑖𝑐𝑎𝑙𝑆𝑦𝑛𝑑𝑟𝑜𝑚𝑒*_ij_* = binary indicator for presence (1) or absence (0) of a specific clinical syndrome; 𝑋_!"_ = vector of covariates (age, sex, illness severity, oedema); *β* = corresponding vector of coefficients for covariates; 𝑀𝑈𝐴𝐶 = mid-upper arm circumference treated as continuous variable to indicate wasting; 𝑀𝑈𝐴𝐶 𝑥 𝐶𝑙𝑖𝑛𝑖𝑐𝑎𝑙𝑆𝑦𝑛𝑑𝑟𝑜𝑚𝑒*_ij_* = interaction term; 𝑏*_oj_* = is a random intercept for site; 𝜖*_ij_* = is the residual error term.

### Multi-omic module preservation and co-regulation

After evaluating the associations between acute illness and biological modules, and assessing the influence of wasting on these relationships, we conducted a module preservation analysis using the *modulePreservation* function from the WGCNA package^75^. This analysis aimed to determine whether the co-expression patterns of individual proteins, metabolites and lipids identified in acutely ill hospitalised children were preserved in well children from similar community settings. Networks from hospitalised children were used as the reference, and those from community children as the test set. Preservation was assessed using two complementary statistics: Zsummary, which combines Zdensity (network density preservation) and Zconnectivity (intramodular connectivity preservation), and medianRank, a size-independent metric useful for comparing preservation across modules. According to standard thresholds, modules with Zsummary < 2 were considered not preserved, 2–10 moderately preserved, and ≥ 10 strongly preserved^75^. Lower medianRank values indicate greater preservation (see Figure 4A–B). Analyses were based on 2,000 permutations.

Following the identification of preserved biological modules shared between acutely ill and well community children, we reconstructed these modules in the well community dataset using the exact feature composition defined in the hospitalised children. For each preserved module, we extracted the corresponding features from well children and applied principal component analysis (PCA) to derive the first principal component (PC1), which was used as the module’s summary measure. Importantly, PC1 is conceptually equivalent to the module eigenfeature computed in WGCNA, which is derived via singular value decomposition and represents the principal axis of variation across all features for a given module. This approach ensured that modules reconstructed in well children reflected the same molecular architecture and summary behaviour as those defined in hospitalised children, enabling biologically consistent and meaningful comparisons of module activity and systems interactions across omic layers as described below.

### Multi-omic integration and network comparisons

Upon the identification of biological modules preserved across acutely ill hospitalised and well community children, we next investigated how these modules interact under conditions of illness versus wellness. To this end, we constructed regularised partial correlation networks using the *EBICglasso* function from the qgraph^76^ R package (v1.9.8), which implements the graphical LASSO algorithm with an extended Bayesian information criterion (EBIC) for model selection. Networks were generated from a subset of strongly and moderately preserved modules, integrating proteomic, metabolomic, lipidomic and haematological features. A tuning parameter of 0.25 and an edge inclusion threshold of 0.06 were applied to optimise the trade-off between model sparsity and sensitivity, while ensuring the stability of inferred associations. As EBICglasso estimates sparse inverse covariance matrices, it enables robust inference of conditional dependencies even in datasets with differing sample sizes. This approach therefore facilitated meaningful comparisons of network structure between hospitalised and community children, while minimising potential biases associated with sample size variability.

To evaluate differences in network topology between hospitalised and community groups, we computed node-level centrality metrics using the *centrality_auto* function in qgraph^76^, which includes strength (sum of absolute partial correlations), closeness (average distance to all other nodes) and betweenness (frequency of a node on shortest paths between other nodes) centralities. To assess how biological coordination shifts between the two states, we computed delta centrality metrics by subtracting values in the community network from those in the hospitalised network (e.g., ΔStrength = Strength for hospitalised network – Strength for community network). These delta metrics were then used to identify nodes with illness-specific increases or decreases in network prominence. We further evaluated whether these differences deviated significantly from zero using a Wilcoxon signed-rank test, an approach conceptually aligned with the network comparison test framework^77^. Networks from hospitalised and well community children are displayed in Figure 4C–D. An overall analysis workflow of the present study is shown in Figure S5.

## ADDITIONAL RESOURCES

We developed an interactive Shiny app that enables exploration of the data and can be accessed via https://mudiboevans.shinyapps.io/Biological-Landscape-Multiomics/.

