## Supplemental Information for "Biological landscape of acute illness in children in sub-Saharan Africa and South Asia"

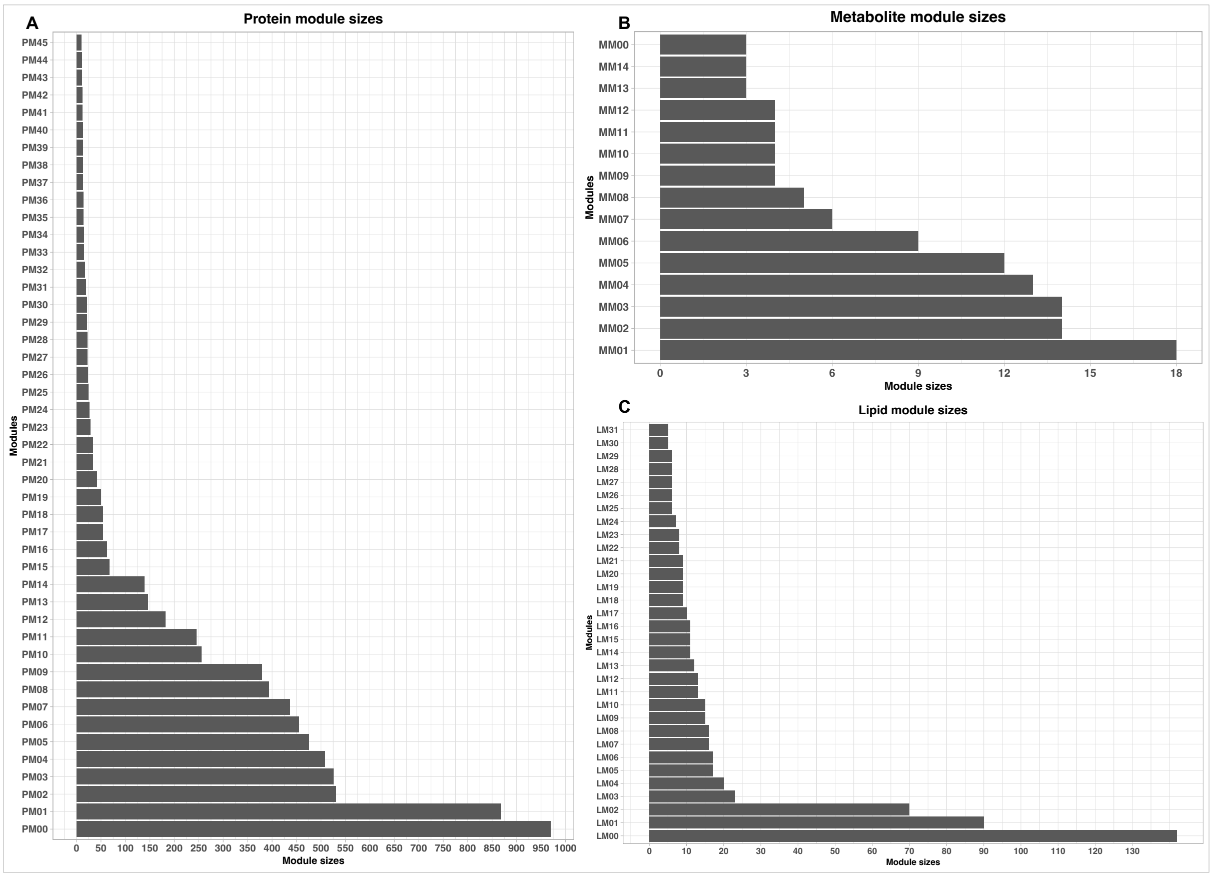

Figure S1. Modules sizes. (A) Number of proteins per module in the protein network. PM00 contain unassigned proteins as such this module was not included in the downstream analyses. Module sizes of assigned proteins ranged between 10 to 868 proteins. (B) Number of metabolites per module in the metabolite network. MM00 contain unassigned metabolites thus this module was not considered in the downstream analyses. The number of metabolites ranged between 3 and 18. (C). Number of lipids per module in the lipid network. LM00 contain unassigned lipids thus this module was not considered for the downstream analyses. The number of lipids ranged between 5 and 90.

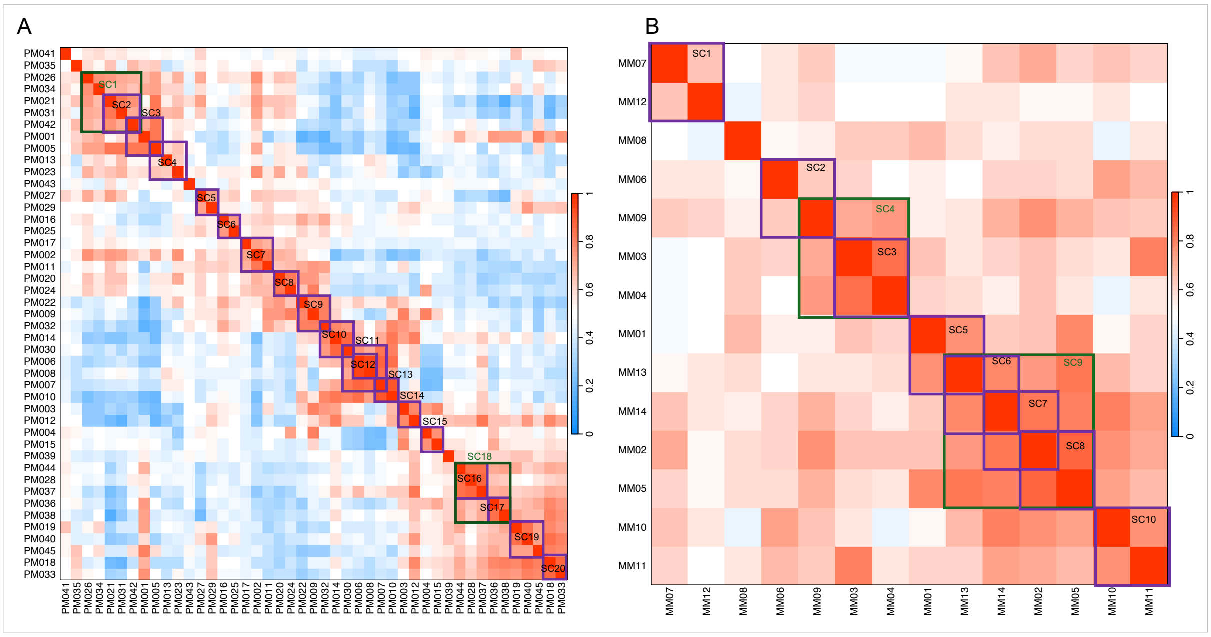

Figure S2. Eigenprotein adjacency heatmap showing the correlation between modules, and superclusters. The highlighted coloured rectangles indicate superclusters with eigenprotein correlations of ≥0.5. (A). Protein superclusters (B) metabolite superclusters. Abbreviations: SC = Supper cluster comprising of tightly correlated modules.

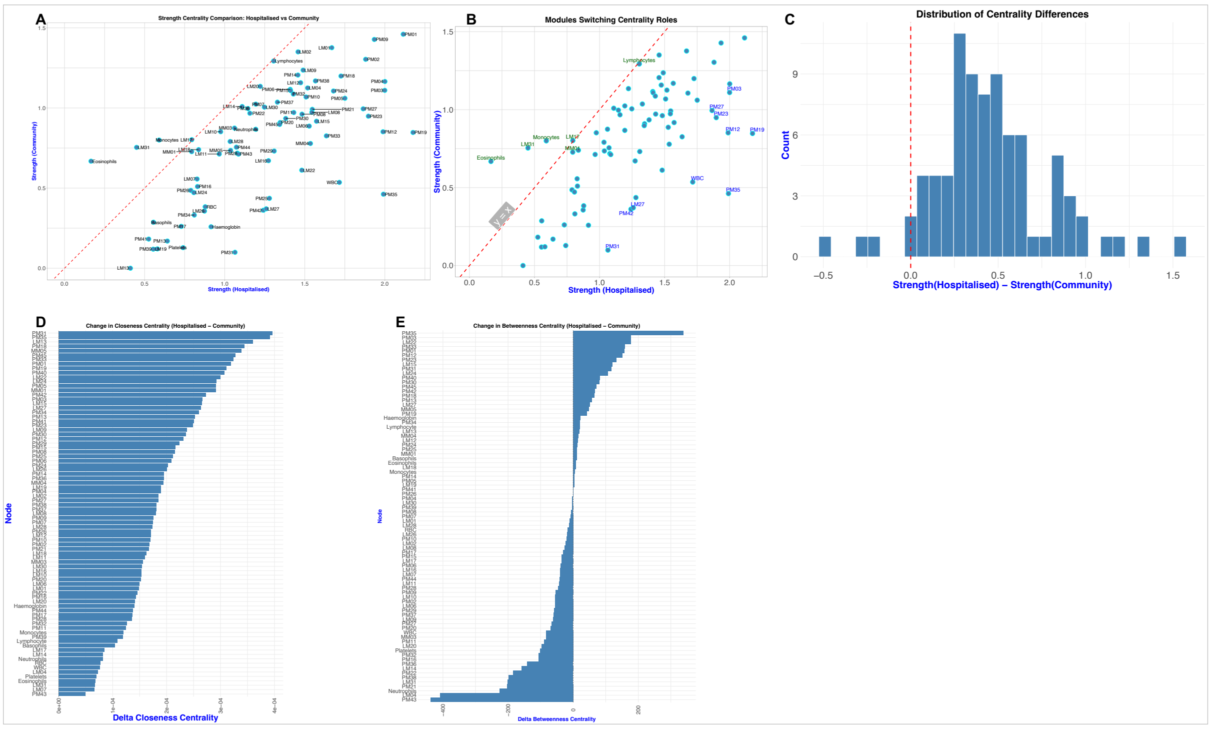

Figure S3. Node-level centrality metrics between the hospitalised and well community children. (A–B). Comparison of node strength centrality between hospitalized and community children. Each point represents a node (e.g., module eigenfeature or hematologic marker), with strength centrality defined as the sum of absolute partial correlation weights connected to that node. The red dashed line represents the identity line ($y = x$), where nodes would have equal centrality in both networks**.** Points below the line are more central in the hospitalized network, whereas points above the line are more central in the community network**.** The large shift of most nodes below the identity line indicates increased system-wide coordination and connectivity during acute illness**.** Strength centrality was estimated using EBIC-regularized partial correlation networks (EBICglasso). (C). Distribution of the delta centrality. (D–E). Closeness and betweenness centrality.

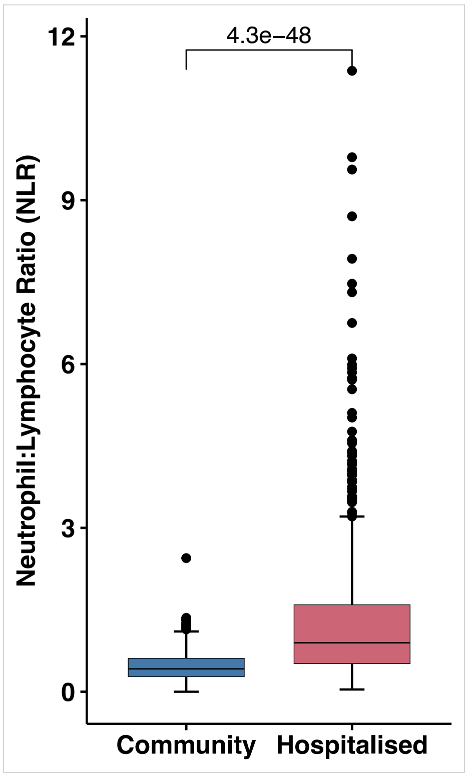

Figure S4. Neutrophil-to-lymphocyte ratio (NLR). Boxplot showing neutrophil-to-lymphocyte ratio between acutely ill hospitalised and well community children. The plot is annotated by p-value following *t.test* indicating significant difference in the mean NLR between the two groups. Boxplot represents the interquartile range (IQR; 25th to 75th percentile), with the median indicated by the centre line. Whiskers extend to 1.5× the IQR and individual points beyond this range represent outliers.

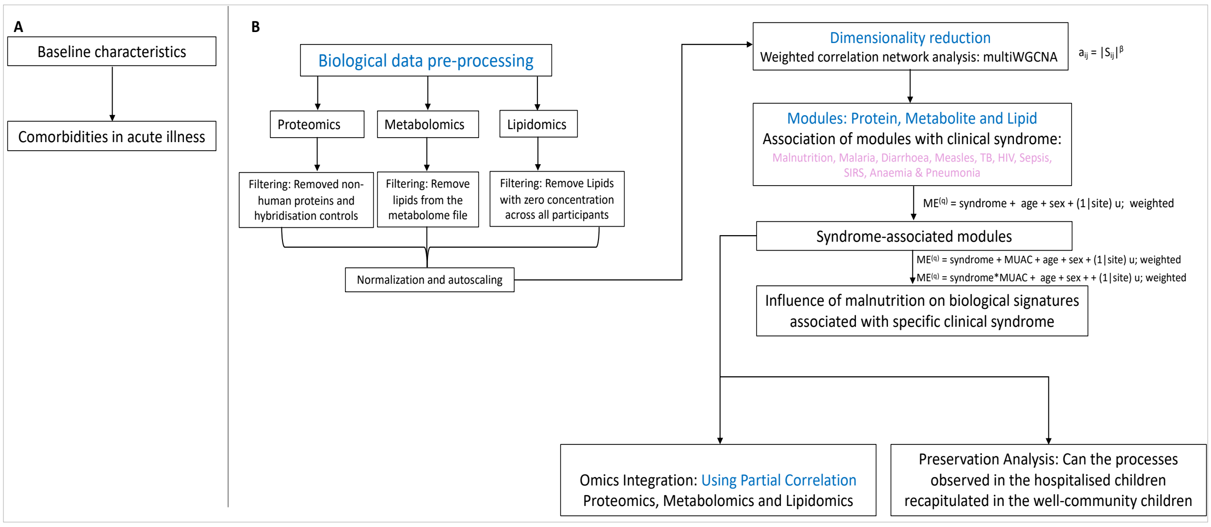

Figure S5. A general analysis workflow of the present study. (A). Analysis of participants characteristics. The figure demonstrates the analyses steps including the profiling of baseline characteristics of the participants. (B). Analysis of biological data including proteomics, metabolomics and lipidomics: pre-processing, dimensionality reduction through weighted correlation network analysis (construction of the network), identification of individual biological modules and their association with clinical syndromes, influence of malnutrition on syndrome-module associations and preservation analysis.

| Table S1. Study participants characteristics at hospital admission by anthropometric categories | | | | | | |
|---|---|---|---|---|---|---|
| **Characteristics at admission** | | | **No wasting** (N = 297) | **Moderate wasting** (N = 230) | **Severe wasting or kwashiorkor** (N = 481) | **All participants** (N=1008) |
| Demographics | | | | | | |
| Sex - Male, n(%) | | | 181 (60.9%) | 144 (62.6%) | 246 (51.1%) | 571 (56.6%) |
| Age, months - Median (IQR) | | | 11.0 (6.8, 16.2) | 10.5 (6.8, 15.3) | 10.3 (6.3, 15.8) | 10.6 (6.5, 15.8) |
| Site N(%) | Kilifi | Kenya | 31 (10.4%) | 15 (6.5%) | 28 (5.8%) | 74 (7.3%) |
| | Migori | | 35 (11.8%) | 24 (10.4%) | 66 (13.7%) | 125 (12.4%) |
| | Nairobi | | 23 (7.7%) | 31 (13.5%) | 48 (10.0%) | 102 (10.1%) |
| | Kampala | Uganda | 32 (10.8%) | 31 (13.5%) | 92 (19.1%) | 155 (15.4%) |
| | Blantyre | Malawi | 42 (14.1%) | 17 (7.4%) | 54 (11.2%) | 113 (11.2%) |
| | Banfora | Burkina Faso | 44 (14.8%) | 32 (13.9%) | 66 (13.7%) | 142 (14.1%) |
| | Karachi | Pakistan | 35 (11.8%) | 22 (9.6%) | 53 (11.0%) | 110 (10.9%) |
| | Dhaka | Bangladesh | 32 (10.8%) | 28 (12.2%) | 48 (10.0%) | 108 (10.7%) |
| | Matlab | | 23 (7.7%) | 30 (13.0%) | 26 (5.4%) | 79 (7.8%) |
| Anthropometric indices – Median (IQR) | | | | | | |
| MUAC | | | 13.45 (12.95, 14.10) | 12.00 (11.60, 12.25) | 10.45 (9.50, 11.25) | 11.75 (10.50, 12.83) |
| WAZ | | | -1.17 (-1.79, -0.43) | -2.58 (-3.29 to -1.94) | -4.18 (-5.16 to -3.36) | -2.87 (-4.17 to -1.58) |
| WHZ | | | -0.82 (-1.45, 0.06) | -2.30 (-2.90 to -1.56) | -3.36 (-4.28 to -2.30) | -2.28 (-3.43 to -1.08) |
| HAZ | | | -1.17 (-1.92, -0.15) | -1.81 (-2.61 to -1.01) | -3.17 (-4.16 to -2.17) | -2.15 (-3.30 to -1.15) |
| Oedema | | | 0 (0.0%) | 0 (0.0%) | 139 (28.9%) | 139 (13.8%) |
| Clinical illness at admission | | | | | | |
| Diarrhoea | | | 139 (47%) | 134 (58%) | 280 (58%) | 553 (55%) |
| Malaria Positive (RDT) | | | 53 (19%) | 27 (14%) | 28 (7.7%) | 108 (13%) |
| Measles | | | 9 (3.3%) | 9 (4.6%) | 6 (1.7%) | 24 (2.9%) |
| Pulmonary TB | | | 2 (0.7%) | 4 (2.0%) | 15 (4.1%) | 21 (2.5%) |
| Pneumonia[a] | | | 129 (43%) | 98 (43%) | 172 (36%) | 399 (40%) |
| Sepsis | | | 39 (13%) | 29 (13%) | 95 (20%) | 163 (16%) |
| SIRS[b] | | | 114 (38%) | 93 (40%) | 182 (38%) | 389 (39%) |
| HIV status | | | | | | |
| HUU | | | 280 (94%) | 205 (89%) | 373 (78%) | 858 (85%) |
| HEU | | | 13 (4.4%) | 15 (6.5%) | 46 (9.6%) | 74 (7.3%) |
| HIV+ | | | 4 (1.3%) | 10 (4.3%) | 62 (13%) | 76 (7.5%) |
| Anaemia[c] | | | | | | |
| None | | | 56 (19%) | 34 (15%) | 89 (19%) | 179 (18%) |
| Mild | | | 74 (25%) | 48 (21%) | 91 (19%) | 213 (21%) |
| Moderate | | | 133 (45%) | 109 (47%) | 222 (46%) | 464 (46%) |
| Severe | | | 34 (11%) | 39 (17%) | 79 (16%) | 152 (15%) |
| Signs of shock[d] | | | | | | |
| None | | | 168 (57%) | 136 (59%) | 266 (55%) | 570 (57%) |
| Some (≥1) | | | 129 (43%) | 94 (41%) | 215 (45%) | 438 (43%) |
| Illness severity signs at admission | | | | | | |
| Low | | | 40 (13%) | 38 (17%) | 100 (21%) | 178 (18%) |
| Medium | | | 118 (40%) | 68 (30%) | 148 (31%) | 334 (33%) |
| High | | | 139 (47%) | 124 (54%) | 233 (48%) | 496 (49%) |
| Haematology – Median (IQR) | | | | | | |
| Haemoglobin, g/dL | | | 9.70 (8.40, 10.90) | 9.50 (8.20, 10.70) | 9.40 (7.90, 10.70) | 9.50 (8.10, 10.80) |
| RBC, $x10^6/\mu L$ | | | 4.60 (4.01, 5.01) | 4.45 (3.72, 5.01) | 4.13 (3.40, 4.72) | 4.35 (3.59, 4.87) |
| WBC, $x10^3/\mu L$ | | | 12 (9, 18) | 13 (10, 19) | 13 (10, 18) | 13 (9, 18) |
| Platelets, $x10^3/\mu L$ | | | 386 (250, 503) | 423 (261, 554) | 359 (195, 546) | 383 (220, 536) |
| Neutrophils, $x10^3/\mu L$ | | | 5.2 (3.5, 8.3) | 5.7 (3.4, 9.5) | 5.0 (3.0, 8.3) | 5.3 (3.3, 8.6) |
| Lymphocytes, $x10^3/\mu L$ | | | 4.9 (3.4, 7.9) | 5.7 (3.8, 8.4) | 6.0 (3.8, 8.3) | 5.6 (3.6, 8.2) |
| Monocytes, $x10^3/\mu L$ | | | 0.91 (0.53, 1.60) | 0.98 (0.60, 1.59) | 1.02 (0.65, 1.71) | 0.98 (0.60, 1.66) |

| | | | | |
|---|---|---|---|---|
| Eosinophils, x$10^3$/μL | 0.06 (0.01, 0.15) | 0.05 (0.01, 0.19) | 0.06 (0.01, 0.15) | 0.06 (0.01, 0.15) |
| Basophils, x$10^3$/μL | 0.06 (0.02, 0.14) | 0.05 (0.02, 0.16) | 0.06 (0.02, 0.16) | 0.05 (0.02, 0.15) |
| Biochemistry – Median (IQR) | | | | |
| Sodium, IU/L | 135 (133, 139) | 135 (132, 139) | 134 (130, 137) | 135 (131, 138) |
| Potassium, IU/L | 4.20 (3.70, 4.80) | 4.20 (3.58, 4.77) | 3.90 (3.03, 4.80) | 4.10 (3.40, 4.80) |
| Calcium, Mmol/L | 2.39 (2.25, 2.54) | 2.36 (2.21, 2.52) | 2.23 (2.00, 2.49) | 2.31 (2.13, 2.51) |
| Magnesium, Mmol/L | 0.96 (0.88, 1.07) | 0.96 (0.87, 1.08) | 0.95 (0.84, 1.12) | 0.96 (0.86, 1.10) |
| Urea, Mmol/L | 3.3 (2.1, 5.8) | 3.0 (1.9, 7.0) | 3.3 (1.9, 7.0) | 3.3 (2.0, 6.5) |
| Creatinine, μmol/L | 22 (2, 29) | 20 (7, 31) | 20 (0, 35) | 21 (0, 31) |
| Albumin, g/L | 40 (34, 43) | 40 (36, 43) | 32 (19, 40) | 38 (26, 42) |
| Bilirubin, μmol/μL | 4 (0, 6) | 4 (2, 8) | 3 (1, 7) | 4 (1, 7) |
| Ionised phosphate, IU/L | 1.65 (1.38, 2.28) | 1.60 (1.31, 2.13) | 1.64 (1.20, 3.20) | 1.62 (1.28, 2.90) |
| Alkaline phosphate, IU/L | 211 (163, 279) | 199 (147, 274) | 169 (123, 238) | 188 (137, 265) |

Data are median (IQR) or count, n (%). Abbreviations and definitions: RDT = rapid diagnostic test; TB = tuberculosis; MUAC = mid-upper arm circumference; WAZ = weight-for-age z score; WHZ = weight-for-height z score; HAZ = height-for-age z score; IQR = interquartile range; HUU = HIV unexposed uninfected; HEU = HIV exposed but uninfected; RBC = R red blood cell; WBC = white blood cell; a = SIRS defined based on presence of two of the following four criteria: heart rate low (<90 bpm) or high (>180 bpm), temperature low (<36·0°C) or high (≥38·5°C), respiratory rate high (>34 breaths per min) and white blood cell count low (<5·0 cells per μL) or high (>17·5 cells per μL); b = severe pneumonia defined as cough or difficulty in breathing with oxygen saturation <90%, central cyanosis or grunting; very severe chest indrawing or inability to breastfeed or drink; or lethargy, reduced level of consciousness, or convulsions; c = Anaemia by haemoglobin defined as none (>110 g/L), mild (100–110 g/L), moderate (70–100 g/L) and severe (<70 g/L); d = signs of shock based on capillary refill time >3 s, upper limb temperature gradient, weak pulse.

Table S2. Study participants characteristics at hospital admission based on omics data availability

| Characteristics | | | Proteomics | | Metabolomics | | Lipidomics | |
|---|---|---|---|---|---|---|---|---|
| | | | **With: N = 944** | **Without: N = 64** | **With: N = 955** | **Without: N = 53** | **With: N = 905** | **Without: N = 103** |
| Demographics | | | | | | | | |
| Sex - Male, n(%) | | | 533 (56.5%) | 38 (59.4%) | 540 (56.5%) | 31 (58.5%) | 513 (56.7%) | 58 (56.3%) |
| Age, months - Median (IQR) | | | 10.6 (6.6, 15.8) | 9 (5, 16) | 10.6 (6.6, 15.8) | 9.6 (6.1, 15.5) | 10.6 (6.6, 15.9) | 9.9 (6.4, 15.2) |
| Site N(%) | Kilifi | Kenya | 74 (7.8%) | - | 74 (7.7%) | - | 72 (8.0%) | 2 (1.9%) |
| | Migori | | 114 (12.1%) | 11 (17.2%) | 113 (11.8%) | 12 (22.6%) | 108 (11.9%) | 17 (16.5%) |
| | Nairobi | | 100 (10.6%) | 2 (3.1%) | 99 (10.4%) | 3 (5.7%) | 96 (10.6%) | 6 (5.8%) |
| | Kampala | Uganda | 149 (15.8%) | 6 (9.4%) | 147 (15.4%) | 8 (15.1%) | 124 (13.7%) | 31 (30.1%) |
| | Blantyre | Malawi | 91 (9.6%) | 22 (34.4%) | 96 (10.1%) | 17 (32.1%) | 84 (9.3%) | 29 (28.2%) |
| | Banfora | Burkina Faso | 140 (14.8%) | 2 (3.1%) | 141 (14.8%) | 1 (1.9%) | 137 (15.1%) | 5 (4.9%) |
| | Karachi | Pakistan | 91 (9.6%) | 19 (29.7%) | 98 (10.3%) | 12 (22.6%) | 98 (10.8%) | 12 (11.7%) |
| | Dhaka | Bangladesh | 107 (11.3%) | 1 (1.6%) | 108 (11.3%) | - | 108 (11.9%) | - |
| | Matlab | | 78 (8.3%) | 1 (1.6%) | 79 (8.3%) | - | 78 (8.6%) | 1 (1.0%) |
| Anthropometric indices – Median (IQR) | | | | | | | | |
| MUAC | | | 11.80 (10.55, 12.88) | 11.50 (10.35, 12.18) | 11.80 (10.50, 12.80) | 11.50 (10.55, 12.95) | 11.80 (10.55, 12.85) | 11.50 (10.40, 12.75) |
| WAZ | | | -2.84 (-4.15, -1.56) | -3.47 (-4.82 to-1.82) | -2.84 (-4.17 to -1.59) | -3.49 (-4.19 to -1.57) | -2.83 (-4.15 to -1.58) | -3.45 (-4.31 to -1.57) |
| WHZ | | | -2.30 (-3.44, -1.09) | -1.98 (-3.08 to -0.71) | -2.30 (-3.42 to -1.10) | -2.14 (-3.52 to -0.47) | -2.29 (-3.42 to -1.10) | -2.20 (-3.44 to -0.85) |
| HAZ | | | -2.10 (-3.26, -1.10) | -2.72 (-4.39 to -1.55) | -2.12 (-3.29 to -1.13) | -2.64 (-4.28 to -1.32) | -2.11 (-3.26 to -1.10) | -2.64 (-4.02 to -1.29) |
| Anthropometric status | | | | | | | | |
| NW | | | 285 (30%) | 12 (19%) | 281 (29%) | 16 (30%) | 268 (30%) | 29 (28%) |
| MW | | | 213 (23%) | 17 (27%) | 221 (23%) | 9 (17%) | 213 (24%) | 17 (17%) |
| SW | | | 446 (47%) | 35 (55%) | 453 (47%) | 28 (53%) | 424 (47%) | 57 (55%) |
| Oedema | | | 128 (13.6%) | 11 (17.2%) | 130 (13.6%) | 9 (17.0%) | 118 (13.0%) | 21 (20.4%) |
| Clinical illness at admission | | | | | | | | |
| Diarrhoea | | | 520 (55%) | 33 (52%) | 524 (55%) | 29 (55%) | 499 (55%) | 54 (52%) |
| Malaria Positive (RDT) | | | 100 (13%) | 8 (15%) | 105 (13%) | 3 (7.7%) | 100 (13%) | 8 (11%) |
| Measles | | | 24 (3.1%) | 54 (100%) | 24 (3.0%) | 39 (100%) | 24 (3.2%) | 75 (100%) |
| Pulmonary tuberculosis (TB) | | | 20 (2.6%) | 1 (1.9%) | 21 (2.6%) | | 17 (2.2%) | 4 (5.3%) |
| Pneumonia[a] | | | 380 (40%) | 19 (30%) | 383 (40%) | 16 (30%) | 363 (40%) | 36 (35%) |
| Sepsis | | | 150 (16%) | 13 (20%) | 152 (16%) | 11 (21%) | 138 (15%) | 25 (24%) |
| SIRS[b] | | | 364 (39%) | 25 (39%) | 373 (39%) | 16 (30%) | 353 (39%) | 36 (35%) |
| HIV status | | | | | | | | |
| HUU | | | 812 (86%) | 46 (72%) | 821 (86%) | 37 (70%) | 785 (87%) | 73 (71%) |
| HEU | | | 65 (6.9%) | 9 (14%) | 67 (7.0%) | 7 (13%) | 59 (6.5%) | 15 (15%) |
| HIV+ | | | 67 (7.1%) | 9 (14%) | 67 (7.0%) | 9 (17%) | 61 (6.7%) | 15 (15%) |
| Anaemia[c] | | | | | | | | |
| None | | | 171 (18%) | 8 (13%) | 173 (18%) | 6 (11%) | 161 (18%) | 18 (17%) |
| Mild | | | 200 (21%) | 13 (20%) | 200 (21%) | 13 (25%) | 187 (21%) | 26 (25%) |
| Moderate | | | 426 (45%) | 38 (59%) | 438 (46%) | 26 (49%) | 418 (46%) | 46 (45%) |
| Severe | | | 147 (16%) | 5 (7.8%) | 144 (15%) | 8 (15%) | 139 (15%) | 13 (13%) |
| Signs of shock[d] | | | | | | | | |
| None | | | 527 (56%) | 43 (67%) | 539 (56%) | 31 (58%) | 503 (56%) | 67 (65%) |
| Some (≥1) | | | 417 (44%) | 21 (33%) | 416 (44%) | 22 (42%) | 402 (44%) | 36 (35%) |

| | | | | | | |
|---|---|---|---|---|---|---|
| Signs of illness severity at admission | | | | | | |
| Low | 166 (18%) | 12 (19%) | 170 (18%) | 8 (15%) | 155 (17%) | 23 (22%) |
| Medium | 311 (33%) | 23 (36%) | 311 (33%) | 23 (43%) | 299 (33%) | 35 (34%) |
| High | 467 (49%) | 29 (45%) | 474 (50%) | 22 (42%) | 451 (50%) | 45 (44%) |
| Haematology – Median (IQR) | | | | | | |
| Haemoglobin, g/dL | 9.50 (8.10, 10.80) | 9.60 (8.15, 10.60) | 9.50 (8.10, 10.80) | 9.65 (7.30, 10.60) | 9.50 (8.10, 10.80) | 9.80 (7.90, 11.00) |
| Red blood cell (RBC), x10$^6$/μL | 4.36 (3.59, 4.88) | 4.18 (3.77, 4.67) | 4.34 (3.59, 4.87) | 4.44 (3.77, 4.97) | 4.33 (3.57, 4.87) | 4.50 (3.88, 5.10) |
| White blood cell (WBC), x10$^3$/μL | 13 (9, 18) | 15.7 (8.5, 18.7) | 13 (9, 18) | 12.7 (9.9, 17.8) | 13 (9, 18) | 13 (10, 18) |
| Platelets, x10$^3$/μL | 382 (219, 533) | 395 (258, 590) | 382 (217, 533) | 437 (265, 586) | 382 (223, 534) | 427 (187, 569) |
| Neutrophils, x10$^3$/μL | 5.3 (3.3, 8.4) | 6.6 (2.5, 11.5) | 5.3 (3.3, 8.6) | 5.4 (3.9, 8.7) | 5.3 (3.3, 8.6) | 4.5 (3.0, 6.8) |
| Lymphocytes, x10$^3$/μL | 5.6 (3.6, 8.2) | 4.4 (3.3, 7.1) | 5.6 (3.6, 8.2) | 4.65 (2.68, 7.12) | 5.5 (3.6, 8.1) | 6.8 (4.4, 8.9) |
| Monocytes, x10$^3$/μL | 0.98 (0.60, 1.64) | 1.1 (0.7, 2.2) | 0.98 (0.60, 1.64) | 1.47 (0.90, 2.56) | 0.98 (0.60, 1.64) | 1.04 (0.62, 1.82) |
| Eosinophils, x10$^3$/μL | 0.06 (0.01, 0.15) | 0.10 (0.04, 0.22) | 0.06 (0.01, 0.15) | 0.08 (0.05, 0.15) | 0.05 (0.01, 0.15) | 0.09 (0.04, 0.15) |
| Basophils, x10$^3$/μL | 0.05 (0.02, 0.15) | 0.10 (0.00, 0.21) | 0.05 (0.02, 0.15) | 0.10 (0.00, 0.21) | 0.05 (0.02, 0.14) | 0.14 (0.05, 0.26) |
| Biochemistry – Median (IQR) | | | | | | |
| Sodium, IU/L | 135 (131, 138) | 137 (132, 139) | 135 (131, 138) | 136 (128, 138) | 135 (131, 138) | 136 (130, 139) |
| Potassium, IU/L | 4.10 (3.37, 4.71) | 4.70 (3.87, 5.20) | 4.10 (3.40, 4.72) | 4.75 (3.80, 5.30) | 4.10 (3.39, 4.70) | 4.63 (3.75, 5.15) |
| Calcium, Mmol/L | 2.30 (2.12, 2.49) | 2.64 (2.30, 8.40) | 2.31 (2.12, 2.50) | 2.62 (2.22, 8.40) | 2.31 (2.13, 2.50) | 2.34 (2.15, 2.60) |
| Magnesium, Mmol/L | 0.96 (0.86, 1.09) | 1.11 (0.87, 2.12) | 0.96 (0.86, 1.09) | 1.20 (0.92, 2.20) | 0.96 (0.86, 1.10) | 0.95 (0.82, 1.10) |
| Urea, Mmol/L | 3.2 (1.9, 6.3) | 5 (3, 10) | 3.2 (1.9, 6.4) | 6 (3, 11) | 3.2 (1.9, 6.3) | 5 (2, 9) |
| Creatinine, μmol/L | 21 (1, 32) | 1 (0, 26) | 21 (1, 32) | 0 (0, 20) | 21 (1, 32) | 0 (0, 22) |
| Albumin, g/L | 38 (26, 42) | 34 (4, 43) | 38 (26, 42) | 34 (5, 40) | 38 (26, 42) | 35 (24, 43) |
| Bilirubin, μmol/μL | 4 (1, 7) | 1 (0, 9) | 4 (1, 7) | 1 (0, 6) | 4 (1, 7) | 1 (0, 5) |
| Ionised phosphate, IU/L | 1.62 (1.28, 2.72) | 1.97 (1.33, 4.90) | 1.61 (1.27, 2.72) | 3.53 (1.71, 5.25) | 1.61 (1.26, 2.56) | 3.12 (1.55, 4.56) |
| Alkaline phosphate, IU/L | 188 (138, 264) | 182 (133, 302) | 187 (137, 263) | 196 (138, 330) | 188 (138, 264) | 187 (135, 284) |

Data are median (IQR) or count, n (%). Abbreviations and definitions: RDT = rapid diagnostic test; IQR = interquartile range; a = SIRS defined based on presence of two of the following four criteria: heart rate low (<90 bpm) or high (>180 bpm), temperature low (<36·0°C) or high (≥38·5°C), respiratory rate high (>34 breaths per min) and white blood cell count low (<5·0 cells per μL) or high (>17·5 cells per μL); b = pneumonia defined as cough or difficulty in breathing with oxygen saturation <90%, central cyanosis or grunting; very severe chest indrawing or inability to breastfeed or drink; or lethargy, reduced level of consciousness, or convulsions; c = Anaemia by haemoglobin defined as none (>110 g/L), mild (100–110 g/L), moderate (70–100 g/L) and severe (<70 g/L); d = signs of shock based on capillary refill time >3 s, upper limb temperature gradient, weak pulse.

Table S3. Significantly over-represented biological pathways

| Module | Size | Enriched Pathways | Adjusted P-value | Database |
|---|---|---|---|---|
| PM01 | 868 | Vascular endothelial growth factor receptor-1 signaling pathway | 0.0030 | DAVID |
| | | Angiogenesis<br>Axonogenesis | 2.2800e-16<br>1.5700e-29 | String |
| | | Axonogenesis | 1.4320e-62 | WebGestalt |
| PM02 | 530 | Innate immune response | 0.0231 | DAVID |
| | | Th1 and Th2 cell differentiation | 0.0224 | String |
| | | Receptor signaling pathway via STAT | 0.0045 | WebGestalt |
| PM03 | 525 | Telomere organization<br>Epigenetic regulation of gene expression | 6.2450e-05<br>4.7820e-04 | DAVID |
| | | Regulation of protein phosphorylation | 6.3100e-18 | String |
| | | Regulation of lymphocyte activation | 8.8630e-9 | WebGestalt |
| PM04 | 508 | Defense response to Gram-negative bacterium | 0.0283 | DAVID |
| | | Defense response to bacterium | 0.0491 | String |
| | | Defense response to bacterium | 0.0030 | WebGestalt |
| PM05 | 475 | Collagen fibril organization<br>Extracellular matrix organization | 5.2760e-03<br>2.1470e-07 | DAVID |
| | | Extracellular matrix organization<br>Ossification | 2.0200e-07<br>1.1700e-06 | String |
| | | Extracellular structure organization | 2.9530e-10 | WebGestalt |
| PM06 | 455 | Regulation of protein localization to plasma membrane | 4.4923e-03 | DAVID |
| | | Intracellular protein transport | 9.9200e-13 | String |
| | | Establishment of protein localization | 1.1708e-10 | WebGestalt |
| PM07 | 436 | ESCRT III complex disassembly<br>Purine nucleotide biosynthetic process | 2.0992e-03<br>0.0013 | DAVID |
| | | Proteolysis involved in protein catabolic process<br>Nucleotide metabolic process | 3.9100e-11<br>5.0700e-11 | String |
| | | Protein catabolic process | 2.3457e-12 | WebGestalt |
| PM08 | 393 | Regulation of platelet activation<br>Actin filament organization | 8.8358e-10<br>2.3279e-04 | DAVID |
| | | Regulation of platelet activation<br>Actin cytoskeleton organization | 1.8100e-08<br>2.1700e-13 | String |
| | | Platelet activation | 2.2217e-13 | WebGestalt |
| PM09 | 379 | Biological process involved in interspecies interaction between organisms<br>Innate immune response | 0.0259<br>0.0355 | String |
| | | Biological process involved in interspecies interaction between organisms | 0.0201 | WebGestalt |
| PM10 | 255 | Positive regulation of cell population proliferation | 0.0000 | DAVID |
| | | Positive regulation of cell population proliferation | 0.0006 | String |
| | | Nucleotide metabolic process | 0.0000 | WebGestalt |
| PM11 | 245 | Positive regulation of fat cell differentiation<br>Inflammatory response | 7.8290e-04<br>1.1570e-02 | DAVID |
| | | Defense response to other organism<br>Positive regulation of fat cell differentiation | 6.0100e-05<br>0.0020 | String |
| | | Defense response to other organism | 0.0000 | WebGestalt |
| PM12 | 182 | Protein refolding<br>Cellular response to heat | 0.0001<br>0.0174 | DAVID |
| | | Protein refolding<br>Response to unfolded protein | 0.0019<br>0.0028 | String |
| | | Protein refolding | 0.0011 | WebGestalt |
| PM13 | 146 | Chemokine-mediated signaling pathway<br>Neutrophil chemotaxis | 2.1077e-02<br>9.6892e-06 | DAVID |
| | | Chemokine-mediated signaling pathway<br>Neutrophil chemotaxis | 5.2800e-07<br>1.0000e-06 | String |
| | | Myeloid leukocyte migration | 8.8099e-12 | WebGestalt |
| PM14 | 139 | mRNA splicing, via spliceosome; mRNA processing | 1.3274e-09 | DAVID |
| | | RNA splicing; mRNA metabolic process | 1.7800e-14 | String |
| | | mRNA metabolic process | 1.2636e-17 | WebGestalt |
| PM15 | 67 | Positive regulation of natural killer cell cytokine production<br>Cytokine-mediated signaling pathway | 0.0451 | DAVID |
| | | Fc-gamma receptor signaling pathway involved in phagocytosis<br>Positive regulation of ERK1 and ERK2 cascade | 0.0313<br>0.0111 | String |
| | | Positive regulation of ERK1 and ERK2 cascade | 0.0093 | WebGestalt |
| PM16 | 62 | Response to oxidative stress | Based on hub protein | |
| PM17 | 54 | Complement activation* | 0.0014 | DAVID |
| PM18 | 54 | Receptor-mediated endocytosis | 0.0123 | DAVID |
| | | Receptor-mediated endocytosis | 0.0007 | String |
| | | Adaptive immune response<br>Cell killing | 0.0000<br>0.0000 | WebGestalt |

| | | | | |
|---|---|---|---|---|
| PM19 | 50 | Antimicrobial humoral immune response mediated by antimicrobial peptide<br>Feeding behaviour | 0.0273<br>0.0022 | DAVID |
| | | Cell wall disruption in another organism<br>Feeding behaviour | 6.5100e-06<br>4.7700e-05 | String |
| | | Defense response to bacterium<br>Feeding behavior | 1.4894e-7 | WebGestalt |
| PM20 | 41 | Phosphatidylethanolamine and phosphatidylcholine biosynthesis | Based on hub protein | |
| PM21 | 33 | Fibrinolysis | 0.0002 | DAVID/String |
| PM22 | 33 | Protein autoubiquitination | 0.0110 | WebGestalt |
| PM23 | 28 | Intraflagellar protein transport | Based on hub protein | |
| PM24 | 26 | Plays a role in adipocyte function and systemic glucose homeostasis | Based on hub protein | |
| PM25 | 24 | Protein-containing complex remodeling | 0.0152 | WebGestalt |
| PM26 | 23 | Chromatin complex remodelling | Based on hub protein | |
| PM27 | 22 | Blood coagulation | 0.0096 | DAVID |
| | | Blood coagulation, fibrin clot formation | 0.0343 | String |
| PM28 | 22 | Alcohol metabolic process<br>Retinol metabolic process | 0.0038<br>0.0326 | DAVID |
| | | Alpha-amino acid metabolic process<br>Carboxylic acid metabolic process | 3.0200e-11<br>8.1900e-18 | String |
| | | Alcohol metabolic process | 8.7831e-10 | WebGestalt |
| PM29 | 21 | Acute-phase response | 0.0016 | DAVID |
| | | Acute-phase response | 0.0019 | String |
| | | Acute inflammatory response | 0.0064 | WebGestalt |
| PM30 | 21 | Cytoplasmic translation | 1.9775e-12 | DAVID |
| | | Cytoplasmic translation | 7.1700e-13 | String |
| | | Cytoplasmic translation | 8.9597e-13 | WebGestalt |
| PM31 | 19 | Zymogen activation | 0.0142 | DAVID |
| PM32 | 17 | ITM2A is involved in activation of T-cells in the immune system and in myocyte differentiation | Based on hub protein | |
| PM33 | 15 | Cellular response protein to viral infection | Based on hub protein | |
| PM34 | 15 | Fibrinolysis* | 0.0156 | DAVID |
| PM35 | 14 | Cell killing | 2.3961e-11 | WebGestalt |
| PM36 | 14 | Axonogenesis<br>Regulation of synapse assembly | 0.0457<br>5.4200e-05 | DAVID<br>String |
| | | Regulation of synapse structure or activity | 0.0004 | WebGestalt |
| PM37 | 13 | Pyruvate metabolic process | 0.0066 | DAVID |
| | | Pyruvate metabolic process | 0.0276 | String |
| | | Small molecule catabolic process | 0.0084 | WebGestalt |
| PM38 | 13 | Involved in cell growth, development and differentiation | Based on hub protein | |
| PM39 | 13 | Muscle contraction<br>Sarcomere organization | 0.0043<br>0.0092 | String |
| | | Muscle contraction<br>Striated muscle cell development | 0.0246 | WebGestalt |
| PM40 | 13 | Disruption of plasma membrane integrity in another organism<br>Défense response to fungus; Innate immune response in mucosa | 0.0004<br>2.6681e-03 | DAVID |
| | | Immune response: Antimicrobial proteins that kills intracellular pathogens | 0.0092 | String |
| | | Disruption of plasma membrane integrity in another organism | 0.0119 | WebGestalt |
| PM41 | 12 | Digestion<br>Proteolysis and lipid catabolism | 4.2706e-08 | DAVID |
| | | Digestion<br>Proteolysis | 2.0800e-07 | String |
| | | Digestion | 0.00000 | WebGestalt |
| PM42 | 12 | Intein-mediated protein splicing<br>Regulation of smoothened signaling pathway | 0.0174<br>0.0370 | String |
| | | Regulation of smoothened signaling pathway | 0.0224 | WebGestalt |
| PM43 | 11 | Protein quality control system which ensures correct protein folding, re-folding of misfolded proteins and control of targeted proteins for degradation | Based on hub protein | |
| PM44 | 11 | Gluconeogenesis<br>NADH oxidation | 0.0063<br>0.0313 | DAVID |
| | | NADH oxidation | 0.0113 | WebGestalt |
| PM45 | 10 | Leukocyte activation<br>T cell activation | 0.0019<br>0.0019 | String |
| | | Leukocyte activation/T cell activation | 0.0375 | WebGestalt |
| SC1-PM26, PM34, PM21, PM31, PM42 | 102 | Negative regulation of fibrinolysis<br>Blood coagulation | 0.0156<br>0.0002 | DAVID |
| | | Blood coagulation<br>Fibrinolysis | 0.0002<br>0.0071 | String |
| | | Blood coagulation | 0.0000 | WebGestalt |

| | | | | |
|---|---|---|---|---|
| SC2-<br>PM21,<br>PM31,<br>PM42 | 64 | Fibrinolysis<br>Blood coagulation | 0.0076<br>0.0029 | DAVID |
| | | Fibrinolysis<br>Blood coagulation<br>Negative regulation of proteolysis | 0.0028<br>0.0150<br>0.0084 | String |
| | | Negative regulation of fibrinolysis | 0.0004 | WebGestalt |
| SC3-<br>PM42,<br>PM01,<br>PM05 | 135<br>5 | Vascular endothelial growth factor receptor-1 signaling pathway<br>Glycosaminoglycan catabolic process<br>Glomerular capillary formation | 8.5314e-23<br>0.0162<br>0.0213 | DAVID |
| | | Angiogenesis | 1.2200e-16 | String |
| | | Axon development | 3.5017e-74 | WebGestalt |
| SC4-<br>PM05,<br>PM13,<br>PM23 | 649 | Neutrophil chemotaxis | 0.0365 | DAVID |
| | | Positive regulation of leukocyte migration | 8.7700e-09 | String |
| | | Chemotaxis | 1.1404e-15 | WebGestalt |
| SC5-<br>PM27,<br>PM29 | 43 | Acute-phase response | 3.5748e-06 | DAVID |
| | | Acute-phase response | 7.2000e-06 | String |
| | | Acute-phase response | 0.0000 | WebGestalt |
| SC6-<br>PM16,<br>PM25 | 86 | Regulation of small molecule metabolic process; Small molecules in GO include monosaccharides but exclude disaccharides and polysaccharides | 0.0197 | String |
| | | Regulation of small molecule metabolic process | 0.0144 | WebGestalt |
| SC7-<br>PM17,<br>PM02,<br>PM11 | 829 | Complement activation,<br>Complement activation, classical pathway | 0.0014<br>0.0047 | DAVID |
| | | Adaptive immune response | 6.9700e-06 | String |
| | | Adaptive immune response | 0.0000 | WebGestalt |
| SC8-<br>PM20,<br>PM24 | 667 | No significantly enriched pathway | ns | DAVID<br>String<br>WebGestalt |
| SC9-<br>PM22,<br>PM09,<br>PM32 | 429 | Biological process involved in interspecies interaction between organisms<br>Regulation of cell death | 0.0073<br>0.0073 | String |
| SC10-<br>PM32,<br>PM14,<br>PM30 | 177 | Cytoplasmic translation<br>mRNA processing | 5.9171e-10<br>3.2377e-07 | DAVID |
| | | RNA splicing, via transesterification reactions<br>Cytoplasmic translation | 1.3500e-12<br>6.4500e-10 | String |
| | | mRNA processing | 4.8725e-15 | WebGestalt |
| SC11-<br>PM30,<br>PM06,<br>PM08,<br>PM07 | 130<br>5 | Regulation of lamellipodium assembly<br>Actin filament depolymerisation | 0.0470<br>0.0004 | DAVID |
| | | Actin cytoskeleton organisation | 1.9600e-16 | String |
| | | Actin cytoskeleton organisation | 1.7747e-11 | WebGestalt |
| SC12-<br>PM06,<br>PM08 | 848 | Regulation of lamellipodium assembly<br>Actin filament depolymerisation | 0.0002<br>0.0086 | DAVID |
| | | Actin cytoskeleton organisation | 5.3000e-19 | String |
| | | Actin filament organization | 3.7013e-16 | WebGestalt |
| SC13-<br>PM07,<br>PM10 | 691 | Viral budding from plasma membrane<br>ESCRT III complex disassembly<br>GMP biosynthetic process | 0.0284<br>0.0050<br>0.0306 | DAVID |
| | | Purine ribonucleoside monophosphate metabolic process | 1.2800e-08 | String |
| | | Ribose phosphate metabolic process | 1.5186e-10 | WebGestalt |
| SC14-<br>PM03,<br>PM12 | 707 | Epigenetic regulation of gene expression<br>Killing of cells of another organism | 0.0086<br>0.0005 | DAVID |
| | | Positive regulation of protein phosphorylation | | String |
| | | Regulation of lymphocyte activation | 2.5671e-9 | WebGestalt |
| SC15-<br>PM04,<br>PM15 | 575 | Defense response to Gram-negative bacterium<br>Cytokine-mediated signaling pathway | 0.0210<br>0.0248 | DAVID |
| | | Cell surface receptor signaling pathway | 2.8400e-09 | String |
| | | Odontogenesis | 0.0003 | WebGestalt |
| SC16-<br>PM44,<br>PM28,<br>PM37 | 46 | NADH oxidation | 0.0105 | DAVID |
| | | Carboxylic acid metabolic process<br>Alpha-amino acid metabolic process | 4.9800e-22<br>1.8700e-09 | String |
| | | Alcohol metabolic process | 4.0474e-15 | WebGestalt |
| SC17-<br>PM36,<br>PM38 | 27 | Neuronal signal transduction | 0.01350 | DAVID |
| | | Neuronal signal transduction | 0.0014 | String |
| | | Regulation of synapse structure or activity | 0.0145 | WebGestalt |
| SC18-<br>PM44,<br>PM28,<br>PM37,<br>PM36,<br>PM38 | 73 | NADH oxidation | 0.0477 | DAVID |
| | | Carboxylic acid metabolic process | 1.1800e-15 | String |
| | | Alcohol metabolic process | 2.1871e-11 | WebGestalt |

| | | | | |
|---|---|---|---|---|
| SC19-PM19, PM40, PM45 | 73 | Disruption of plasma membrane integrity in another organism | 0.0002 | DAVID |
| | | Antimicrobial humoral response<br>Feeding behaviour | 8.5100e-08 0.0003 | String |
| | | Humoral immune response<br>Disruption of anatomical structure in another organism | 2.4622e-8 | WebGestalt |
| SC20-PM18, PM23 | 82 | Receptor-mediated endocytosis<br>Inflammatory response | 0.0359<br>0.0021 | DAVID |
| | | Inflammatory response<br>T-helper 1 type immune response<br>Regulation of adaptive immune response based on somatic | 5.1600e-06<br>0.0004<br>2.0100e-05 | String |
| | | Adaptive immune response<br>Type II interferon production<br>Cell killing | 3.5156e-8 | WebGestalt |
| Homo sapiens used as the reference background for calculating fold enrichment for Gene Ontology enrichment analysis for biological processes. Enrichment was assessed with Fisher's exact or hypergeometric tests, and P value adjusted for Bonferroni correction for DAVID and FDR for WebGestalt and String. *Functional based on the clusters of modules that were tightly correlated with each other forming superclusters. Size column indicates the number of proteins making up each protein module |  |  |  |  |

| Table S4. Metabolite modules and the associated metabolites per module | | | |
|---|---|---|---|
| **Module** | **Size** | **Metabolites** | **Metabolic Function/Pathway** |
| MM01 | 18 | C12:1, C12, C14:2, C14:1, C14, C12DC, C14:2OH, C14:1OH, C16:2, C16:1, C16, C16:2OH, C16:1OH, C16OH, C18:2, C18:1, C18, C18:1OH | Long-chain acylcarnitine metabolism, Mitochondrial β-oxidation, Fatty acid transport and oxidation, Energy metabolism |
| MM02 | 14 | Creatinine, Putrescine, total dimethylarginine, Methylhistidine, Cystathionine, Homocitrulline, Urea, Uric acid, X5-Hydroxy Indoleacetic acid, C4, C3OH, C5:1, C5OH, C9 | Nitrogen metabolism, Urea cycle, Creatine-phosphocreatine system, Muscle metabolism, amino acid metabolism |
| MM03 | 14 | Alanine, Serine, Proline, Threonine, Asparagine, Aspartic acid, Glutamine, Glutamic acid, Methionine, Methionine sulfoxide, Asymmetric dimethylarginine, Lysine, Sarcosine, Hydroxy lysine | Amino acid metabolism (Alanine, Aspartate, and Glutamate metabolism) |
| MM04 | 13 | Valine, Leucine, Isoleucine, alpha-Aminoadipic acid, Phenylalanine, Arginine, Citrulline, Tyrosine, Tryptophan, Kynurenine, Ornithine, Choline, alpha-aminobutyric acid | Branched-chain amino acid metabolism (Valine, Leucine, Isoleucine degradation) |
| MM05 | 12 | gamma-aminobutyric acid, C4:1, C6:1, C6, C5:1DC, C5DC, C8, C5MDC, C7DC, C10:2, C10:1, C10 | Short- and medium-chain acylcarnitine metabolism; Carnitine metabolism, energy production |
| MM06 | 9 | Carnosine, Spermidine, Spermine, Lactic acid, alpha-Ketoglutaric acid,<br>Citric acid, Succinic acid, Fumaric acid, Pyruvic acid | Krebs cycle (TCA cycle), energy production |
| MM07 | 6 | Trimethylamine N-oxide, HPHPA, p-Hydroxyhippuric acid, Hippuric acid, Indole acetic acid, p-Hydroxyphenylacetic acid | Gut microbiota-derived metabolism; Xenobiotic metabolism |
| MM08 | 5 | Taurine, Serotonin, Hypoxanthine, Uridine, Xanthine | Neurotransmitter metabolism |
| MM09 | 4 | Methylmalonic acid, C0, C3, C5 | Vitamin B12 metabolism, propionate metabolism, Odd-chain fatty acid oxidation |
| MM10 | 4 | Betaine, Ethanolamine, N-acetylpetrescine, Homovanillic acid | Choline and Methyl Donor Metabolism, Catecholamine Degradation; Methylation and lipid metabolism |
| MM11 | 4 | Glycine, trans-Hydroxyproline, Histidine, Guanidineacetic acid | Amino acid metabolism, creatine biosynthesis |
| MM12 | 4 | Butyric acid, Propionic acid, Isobutyric acid, Valeric acid | Short-chain fatty acid metabolism; Gut microbiota-derived metabolism |
| MM13 | 3 | beta-Hydroxybutyric acid, C2, C4OH | Ketone body metabolism |
| MM14 | 3 | Creatine, Acetyl-lysine, C3.1 | Creatine phosphate system; Energy metabolism |

Table S5. Lipid modules and the associated lipid per module

| Module | Size | Lipids |
|---|---|---|
| LM01 | 90 | Cer_(42:0-OH), CL_(75:08), CL_(75:07), DG_(32:0), DG_(40:3), DG_(38:3), DG_(42:3), DG_(40:1), Hex-Cer_(34:2-OH), Hex-Cer_(43:1), PC_(39:1), PC_(37:0), PE_(36:3), PE_(34:0), PG_(37:6), PG_(36:4), PG_(40:1), TG_(12:0), TG_(36:0), TG_(57:10), TG_(42:2), TG_(46:4), TG_(57:9), TG_(48:5), TG_(47:4), TG_(42:1), TG_(44:2), TG_(46:3), TG_(48:4), TG_(43:1), TG_(49:5), TG_(45:2), TG_(58:11), TG_(47:3), TG_(50:5), TG_(42:0), TG_(44:1), TG_(46:2), TG_(49:4), TG_(48:3), TG_(43:0), TG_(59:8), TG_(45:1), TG_(50:4), TG_(47:2), TG_(52:5), TG_(58:9), TG_(49:3), TG_(54:6), TG_(51:4), TG_(44:0), TG_(58:8), TG_(46:1), TG_(48:2), TG_(56:7), TG_(50:3), TG_(45:0), TG_(52:4), TG_(47:1), TG_(54:5), TG_(49:2), TG_(51:3), TG_(55:6), TG_(53:4), TG_(56:6), TG_(46:0), TG_(48:1), TG_(50:2), TG_(52:3), TG_(47:0), TG_(54:4), TG_(49:1), TG_(51:2), TG_(56:5), TG_(53:3), TG_(58:6), TG_(55:4), TG_(50:1), TG_(52:2), TG_(48:0), TG_(54:3), TG_(56:4), TG_(51:1), TG_(53:2), TG_(55:3), TG_(58:5), TG_(50:0), TG_(52:1), TG_(54:2), TG_(51:0) |
| LM02 | 70 | Cer_(34:1), Cer_(35:1), Cer_(36:1), Cer_(36:0), Cer_(37:1), Cer_(40:2), Cer_(38:1), Cer_(39:1), Cer_(41:2), Cer_(40:1), Cer_(42:2), Cer_(40:0), Cer_(41:1), Cer_(43:2), Cer_(41:0), Cer_(42:1), Cer_(42:0), Cer_(43:1), Cer_(44:1), CPE_(36:1), CPE_(38:2), CPE_(37:1), CPE_(42:2), CPE_(40:1), CPE_(42:1), GB3_(40:1), Hex-Cer_(35:2), Hex-Cer_(35:1), Hex-Cer_(35:0-OH), Hex-Cer_(36:1-OH), Hex-Cer_(36:2), Hex-Cer_(36:1), Hex-Cer_(37:1), Hex-Cer_(39:1), PC_(34:4), PC_(32:2), PC_(33:2), PC_(30:0), PC_(35:3), PC_(32:1), PC_(34:2), PC_(36:3), PC_(31:0), PC_(33:1), PC_(35:2), PC_(37:3), PC_(32:0), PC_(34:1), PC_(36:2), PC_(33:0), PC_(35:1), PC_(38:3), PC_(37:2), PC_(34:0), PC_(38:2), PC_(36:1), PC_(40:3), PC_(38:1), PC_(37:1), PC_(40:2), PC_(40:1), PC_(42:2), PC_(42:1), PC_(40:0), PE_(31:0), PE_(39:6), PG_(38:5), PS_(33:1), PS_(35:2), PS_(40:5) |
| LM03 | 23 | CL_(70:07), CL_(72:08), CL_(66:04), CL_(70:06), CL_(72:07), CL_(68:04), CL_(70:05), CL_(66:02), CL_(72:06), CL_(70:04), CL_(66:01), CL_(72:05), CL_(70:03), CL_(72:04), Mono-lyso_CL_(52:05), Mono-lyso_CL_(54:06), Mono-lyso_CL_(52:04), Mono-lyso_CL_(56:06), Mono-lyso_CL_(52:03), Mono-lyso_CL_(54:05), Mono-lyso_CL_(56:05), PG_(34:1), PS_(36:2) |
| LM04 | 20 | Cer_(38:1-OH), Lacto-cer_(34:0-OH), LPC_(16:1), LPC_(17:2), LPC_(17:1), LPE_(17:0), LPE_(22:5), LPI_(18:2), LPI_(20:4), LPI_(16:0), LPI_(20:3), LPI_(18:1), LPI_(20:2), LPI_(17:0), LPI_(18:0), LPI_(20:0), Mono-lyso_CL_(53:00), PC_(36:0), PS_(36:4), PS_(37:5) |
| LM05 | 17 | Cer_(36:2), Cer_(39:0), CL_(73:08), CPE_(34:1), CPE_(37:0), DGDG_(34:1), GM1_(34:1-OH), GM2_(34:1-OH), GM1_(36:1-OH), PC_(29:1), PC_(31:2), PE_(33:0), PS_(35:4), SM_(30:0), S_(36:2-OH), TG_(59:6), TG_(59:5) |
| LM06 | 17 | LPC_(14:0), LPC_(18:2), LPC_(20:5), LPC_(15:0), LPC_(16:0), LPC_(18:1), LPC_(17:0), LPC_(18:0), LPC_(22:4), LPC_(19:0), LPC_(20:1), LPC_(20:0), LPC_(22:3), LPC_(22:0), LPC_(24:0), PC_(42:10), S_(36:1-OH) |
| LM07 | 16 | DGDG_(32:2), DGDG_(36:0), DGDG_(38:0), DG_(33:0), DG_(34:0), DG_(35:0), DG_(36:0), DG_(38:0), TG_(49:0), TG_(52:0), TG_(53:0), TG_(54:0), TG_(55:0), TG_(56:0), TG_(57:0), TG_(58:0) |
| LM08 | 16 | GB4_(34:1), SM_(32:1), SM_(32:0), SM_(34:1-OH), SM_(34:1), SM_(35:2), SM_(36:2), SM_(36:1), SM_(38:2), SM_(38:1), SM_(40:2), SM_(42:2), SM_(40:1), S_(42:2-OH), S_(40:1), TG_(58:10) |
| LM09 | 15 | GD3_(34:1-OH), GD3_(34:1), GD1_(34:1), GD1_(36:2), GD3_(36:1), GD1_(36:1), GD3_(40:1), GD1_(42:2), GD3_(42:1), GM1_(36:1), GM2_(36:1), GM3_(36:1), GM3_(40:1), Lacto-cer_(34:1-OH), PG_(37:1) |
| LM10 | 15 | PI_(32:0), PI_(40:6), PC_(37:6), PC_(38:6), PC_(36:4), PC_(38:5), PC_(39:6), PC_(40:6), PC_(38:4), PC_(40:5), PC_(42:8), PC_(40:4), PC_(42:7), PE_(30:1), PG_(38:1) |
| LM11 | 13 | Lacto-cer_(34:2), Lacto-cer_(34:1), Lacto-cer_(34:0), Lacto-cer_(36:1), Lacto-cer_(38:1), Lacto-cer_(40:2), Lacto-cer_(39:1), Lacto-cer_(40:1), Lacto-cer_(42:2), Lacto-cer_(40:0), Lacto-cer_(42:1), Lacto-cer_(44:2), S_(48:1) |
| LM12 | 13 | Cer_(45:1), Hex-Cer_(42:0-OH), SM_(35:1), SM_(37:2), SM_(37:1), SM_(39:1), SM_(40:0-OH), SM_(41:1), SM_(44:2), SM_(42:1), SM_(43:1), SM_(44:1), S_(33:1) |
| LM13 | 12 | CL_(72:09), CL_(71:08), CL_(74:08), CL_(74:04), GB4_(42:2), GB4_(40:1), GB4_(40:0), GB4_(42:1), GB3_(42:1), PS_(40:8), PS_(38:4), PS_(40:4) |
| LM14 | 11 | DG_(34:2), DG_(34:1), DG_(40:4), DG_(35:1), DG_(36:2), DG_(36:1), DG_(38:2), DG_(37:1), DG_(38:1), PS_(38:1), PS_(38:0) |
| LM15 | 11 | Hex-Cer_(36:0-OH), PE_(36:2), PE_(38:6), PE_(36:4), PE_(34:2), PE_(40:6), PE_(34:1), PE_(40:7), PE_(38:4), PE_(36:1), SM_(36:3) |
| LM16 | 11 | PI_(36:4), PI_(34:2), PI_(37:4), PI_(35:2), PI_(37:3), PI_(38:4), PI_(40:5), PI_(38:3), PI_(37:2), PI_(40:4), PI_(38:2) |
| LM17 | 10 | TG_(56:3), TG_(56:2), TG_(54:1), TG_(58:3), TG_(56:1), TG_(57:2), TG_(58:2), TG_(57:1), TG_(58:1), TG_(60:0) |
| LM18 | 9 | GM3_(38:0), SM_(33:0), SM_(34:0), SM_(36:0), SM_(38:0), SM_(40:0), SM_(41:0), SM_(42:0), S_(36:1) |
| LM19 | 9 | CL_(69:08), GB3_(38:0), PC_(42:9), PS_(37:3), PS_(35:1), PS_(39:4), PS_(35:0), PS_(38:2), PS_(37:1) |
| LM20 | 9 | GM1_(32:1), GM1_(32:0), GM3_(32:1), GM3_(32:0), GM1_(34:1), GM1_(34:0), GM2_(34:1), GM2_(34:0), Lacto-cer_(32:1) |
| LM21 | 9 | DG_(36:3), GD1_(42:1), PI_(33:0), PI_(34:0), PI_(37:0), Mono-lyso_CL_(53:06), SM_(37:0), S_(38:1), S_(38:0) |
| LM22 | 8 | GD1_(36:0), GM3_(33:1), GM3_(33:0), GM3_(34:1), GM3_(34:0), GM3_(35:1), Lacto-cer_(33:1), PG_(39:7) |
| LM23 | 8 | PE_(32:1), PE_(37:4), PE_(35:1), PE_(38:3), PE_(40:5), PE_(40:4), PE_(38:2), PE_(38:1) |
| LM24 | 7 | LPE_(18:2), LPE_(20:5), LPE_(16:0), LPE_(18:1), LPE_(18:0), LPE_(20:3), PG_(40:3) |
| LM25 | 6 | CL_(73:07), CL_(76:08), GM1_(40:1-OH), Mono-lyso_CL_(56:03), PG_(42:8), PG_(40:4) |
| LM26 | 6 | GB3_(34:1), Hex-Cer_(34:1), Hex-Cer_(38:1), Hex-Cer_(40:1), Hex-Cer_(41:1), Hex-Cer_(42:1) |
| LM27 | 6 | PG_(34:0), S_(34:1-OH), S_(34:1), S_(42:2), S_(42:1-OH), S_(42:1) |
| LM28 | 6 | PI_(32:1), PI_(33:1), PI_(34:1), PI_(36:2), PI_(35:1), PI_(36:1) |
| LM29 | 6 | GB3_(34:1-OH), GB3_(34:2), GB3_(40:1-OH), GB3_(42:2-OH), GB3_(42:3), GB3_(42:1-OH) |
| LM30 | 5 | PC_(35:4), PC_(36:5), PC_(37:4), PC_(39:5), SM_(33:1) |
| LM31 | 5 | GM2_(32:1), Lacto-cer_(32:1-OH), LPC_(12:0), S_(32:1), S_(32:0) |

Lipid classes: Cer = Ceramide, CL = Cardiolipin, DG = Diglyceride, Hex-Cer = Hexasolyceramide, PC = Phosphatidylcholine, PE = Phosphatidylethanolamine, PI = Phosphoinositol, CPE = Ceramide PE, TG = Triglyceride, LPC = Lysophosphatidylcholine, GM1/GM2/GM3 = Monosialo-ganglioside, GD1/GD3 = Ganglioside, GB3s = Globosides, LPE = Lysophosphatidylethanolamine, PS = Phosphoserine, SM = Sphingomyelin, S = Sulfatide, Lacto-cer = Lactosylceramide, LPI = Lysophosphoinositol, DGDG = Digalactosyldiaclglycerols

| Table S6. Characteristics of the well community participants | | | |
|---|---|---|---|
| **Characteristics of well community children** | | | **N = 300** |
| Demographics | | | |
| Sex - Male, n(%) | | | 157 (52.3%) |
| Age, months - Median (IQR) | | | 12.2 (7.4, 17.8) |
| Site N(%) | Kilifi | Kenya | 30 (10.0%) |
| | Migori | | 37 (12.3%) |
| | Nairobi | | 30 (10.0%) |
| | Kampala | Uganda | 30 (10.0%) |
| | Blantyre | Malawi | 31 (10.3%) |
| | Banfora | Burkina Faso | 32 (10.7%) |
| | Karachi | Pakistan | 50 (16.7%) |
| | Dhaka | Bangladesh | 30 (10.0%) |
| | Matlab | | 30 (10.0%) |
| Anthropometric indices – Median (IQR) | | | |
| Mid-upper arm circumference (MUAC) | | | 13.55 (12.95, 14.45) |
| Weight-for-age z score (WAZ) | | | -1.03 (-1.81 to -0.18) |
| Weight-for-height z score (WHZ) | | | -0.42 (-1.10 to 0.46) |
| Height-for-age z score (HAZ) | | | -1.42 (-2.27 to -0.76) |
| Data are median (IQR) or count, n (%). IQR = interquartile range | | | |
